# Copy number normalization distinguishes differential signals driven by copy number differences in ATAC-seq and ChIP-seq

**DOI:** 10.1101/2024.04.11.588815

**Authors:** Dingwen Su, Moritz Peters, Volker Soltys, Yingguang Frank Chan

## Abstract

A common objective across ATAC-seq and ChIP-seq analyses is to identify differential signals across contrasted conditions. However, in differential analyses, the impact of copy number variation is often overlooked. Here, we demonstrated copy number differences among samples could drive, if not dominate, differential signals. To address this, we propose a pipeline featuring copy number normalization. By comparing the averaged signal per gene copy, it effectively segregates differential signals driven by copy number differences from other factors. Further applying it to Down syndrome, we unveiled distinct dosage-dependent and -independent changes on chromosome 21. Thus, we recommend normalization as a general approach.

## Background

Recent advances in next-generation sequencing assays have boosted the application of high-throughput methods in research. Combined with rapidly decreasing costs, researchers can now routinely apply functional genomic assays such as Assay for Transposase-Accessible Chromatin with high-throughput sequencing (ATAC-seq) and chromatin immunoprecipitation followed by sequencing (ChIP-seq) to profile alterations in histone modifications, protein-DNA interactions and open chromatin landscapes, thus providing information on gene regulation [1–5]. Changes in them are closely associated with early developmental processes, the initiation, and progression of diseases. In the case of cancer, differential signals can serve as potential biomarkers and even therapeutic targets [6], [7].

A critical step in these studies is to identify differential signals between perturbed and control states, along developmental stages or in response to the titration of specific chemical treatments. After read alignment and filtering, a common workflow of the differential analysis for ATAC-seq and ChIP-seq is to first identify a focal set of regions with enriched signal (peaks), then quantify the signals, normalize them and lastly detect differential signals via statistical tests. It is important to note that the differential results can differ greatly depending on bioinformatic pipelines (e.g., ENCODE pipelines) and specific parameter choices, even from the same set of data, as demonstrated by Reske *et al.* [8–12]. Even when using the same data and pipeline, the outcome of the differential analysis may differ depending on how the variation in the data is attributed.

Variations in signals between contrasted conditions arise from various sources, which is ideally but often not limited to the factor(s) of interest. Correcting biases and attributing the variation to factors accordingly in the differential analysis is crucial for obtaining an accurate set of differential signals. One factor often overlooked in the differential analysis is copy number variation (CNV), including chromosomal aberrations such as aneuploidy. Specifically, since the signal captured by ATAC-seq or ChIP-seq is the sum of signals from all gene copies, variation in the underlying copy number or ploidy can directly impact the aggregated signal and the interpretation of differential signals. As a result, differences in copy numbers between samples may drive the identification of differential signals, even without any changes in local differences in chromatin accessibility or binding.

However, commonly used tools to quantify reads/fragments in ATAC-seq or ChIP-seq peaks like *bedtools*, *deeptools*, *htseq-count,* and *featureCounts* in the first place are not inherently designed to distinguish between background and effective signals. As a result, these tools quantify background signals as effective signals, inflating the detected effective signals [13–16]. A higher copy number in a region will thus inflate the signal to a larger extent, increasing the likelihood of misidentifying this region as exhibiting elevated signals. Other tools like *DiffBind* and *csaw* do offer the option to subtract the signals from the background when quantifying the signals in peaks [17–19]. However, by being copy-number blind, the effects of differences in the copy number between samples remain. Meanwhile, CNV and/or aneuploidy are commonplace in cell lines used in biomedical research. It could arise from cancer, defective developmental defects (e.g., trisomy 13 or 21, the latter commonly known as Down syndrome), the process of establishing cell lines and repeated passages in tissue culture [20–23]. Additionally, many tissues are naturally polyploid, such as the salivary gland in *Drosophila melanogaster*, the liver in humans and the developing root in *Arabidopsis thaliana.* Therefore, to accurately identify the alterations due to the factor(s) of interest in biological processes or diseases, it is necessary to take CNV into account as a source of variation in the differential analysis, especially when CNV and aneuploidy may arise as an artifact of cell immortalization.

To address this, we propose a differential analysis pipeline featuring copy number (CN) normalization and we showcase its application and advantages using two examples in biomedical studies. In the first case, we applied it to ATAC-seq and ChIP-seq data generated from cell lines with complex chromosomal aberrations derived from a Bloom syndrome individual and a healthy donor. Using a conventional copy-number blind pipeline, we noticed heavily skewed differential signals toward the sample with relatively higher copy numbers in the corresponding regions. In this case where CNV is not a central factor of interest, applying our pipeline with CN normalization efficiently distinguishes differential signals driven by copy number variations and those due to the disease. In the second case, we applied it to ATAC-seq data generated from trisomy 21 and euploid cell lines. By combining the results from our pipeline with the common workflow, we were able to distinguish the open chromatin regions on chromosome 21 with dosage effects, compensatory effects and copy number-independent regulatory changes.

## Results

### Copy number variation drives the identification of differential signals in ATAC-seq and ChIP-seq

Bloom syndrome (BS) is a rare autosomal recessive disorder with complex clinical manifestations. It is caused by loss-of-function of mutations in the conserved Bloom syndrome RecQ like helicase, *BLM* gene, on chromosome 15. BLM helicase untangles various aberrant DNA structures and thus plays an essential role in maintaining genome integrity [24, 25]. Clinically, BS individuals have strikingly shorter lifespans, in average 26 years. They have an elevated risk of cancer and are predisposed to developing cancer at an early age [26]. At the molecular level, BS cells display characteristic excessive sister chromatid exchange events and hallmarks of genome instability [26–28]. Identifying molecular alterations in BS helps to better understand the pathogenesis of the disease as well as the function of BLM helicase.

As primary samples from BS individuals are not readily available, we used two fibroblast cell lines derived from a BS individual and a healthy donor (wildtype or WT) and profiled their open chromatin landscape with ATAC-seq and endogenously formed G-quadruplex structures via ChIP-seq [29]. The Coriell repository described these cell lines as displaying different and atypical karyotypes as well as chromosomal rearrangement events. To assess copy number differences between the BS and WT cell lines, we used *CNVkit* and whole-genome sequencing data to estimate the copy number ratio (CNR), the ratio of local copy number in a given region in BS against WT [30]. Besides irregular karyotypes with complex aneuploidy in each cell line, there were widespread copy number differences between the two cell lines (**Fig. 1a**, **Additional file 1**). Excluding sex chromosomes and masked regions in the human reference genome (hg38), the BS sample exhibits relative copy number gains and losses in a total of 1273.1 Mb (47.0%) vs. 1437.9 Mb (53.0%), respectively.

**Fig. 1.**
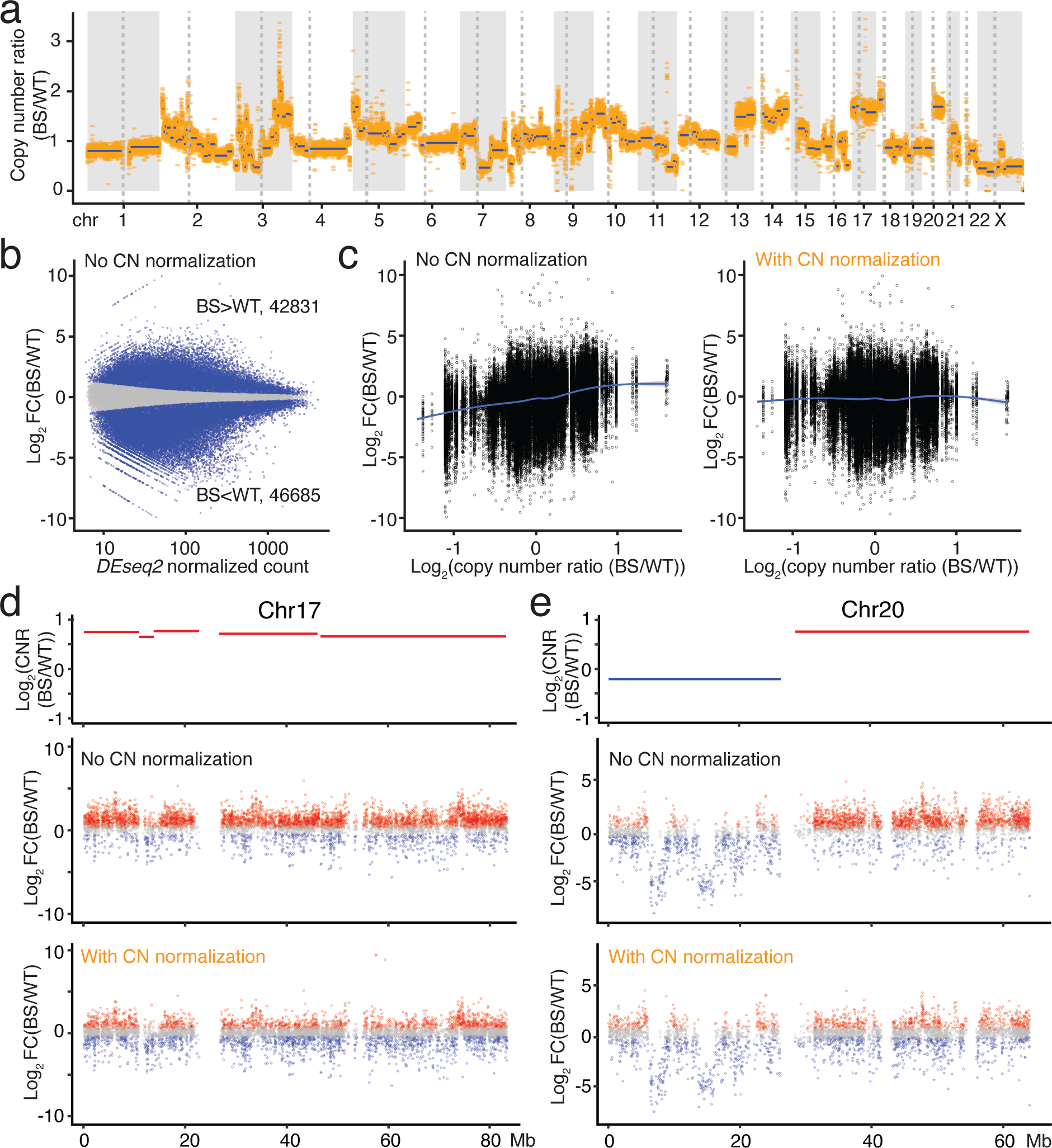
Copy number variation drives the differential signals in ATAC-seq data. **a** Genome-wide copy number ratio (CNR) in the BS sample relative to the WT sample. Orange dots indicate the CNR for each 50kb bin and blue lines indicate DNA segments with the same CNR. **b** MA plot comparing the chromatin accessibility in BS vs. WT samples by *DESeq2*. FC, fold change. **c** The trended biases of differential chromatin accessibility from copy number differences without (left) and with (right) copy number (CN) normalization in ATAC-seq data. Log_2_CNR > 0 and log_2_CNR < 0 imply relative number gain and loss in BS, respectively. The lines represent the locally weighted running line smoother (LOESS) smoothing curve for the data with the grey area indicating the 95% confidence interval for the fitted curve. **d** CNR and the distribution of differential peaks without and with CN normalization on chromosome 17 (chr17). Peaks with adjusted *P* (p-adj) < 0.05 are depicted in blue or red while those with p-adj ≥ 0.05 are shown in grey. **e** CNR and the distribution of differential peaks without and with CN normalization on chr20. Peaks with p-adj < 0.05 are depicted in blue or red while those with p-adj ≥ 0.05 are shown in grey.

To characterize the impacts of BS on chromatin accessibility, we performed differential analysis using a workflow with a few widely used bioinformatic tools. The major steps after read alignment and filtering included: 1) calling peaks with *MACS2*; 2) quantifying signal as the number of reads/fragments in peaks with *htseq-count*; 3) performing data normalization and identifying differential signals with *DESeq2*, specifying genotype ((BS vs. WT) as the contrasting factor [31]. Following this pipeline, we identified 143,460 accessible chromatin regions in WT and BS cells and detected 89,516 (62.40%) significantly differential peaks (herein defined as peaks with adjusted *P* < 0.05) in BS vs. WT. Under this standard pipeline, 42,831 (29.86%) peaks showed increased accessibility, while 46,685 (32.54%) displayed decreased accessibility. When assessing the genome-wide differential signals using an MA plot, we observed no obvious skewness towards the WT or BS sample (**Fig. 1b**). Crucially, we noted a trended CNR-dependent bias in the differential signals (**Fig. 1c**, **left**): there was an over-representation of accessible peaks in regions with copy number gains (log_2_ CNR >0); conversely, less accessible peaks in regions with copy number losses (log_2_ CNR <0).

To better investigate this effect, we examined two chromosomes in detail to see how copy number differences may affect the differential analysis. For chromosome 17 the entire chromosome showed relatively higher copy numbers in BS. About 72% (5239 out of 7231) of the peaks were significantly differential. Notably, there is a very strong directionality to this set: 62.40% (4486) showed increased chromatin accessibility in BS, but only 10.47% (753) with decreased signals. Both of these proportions deviated strongly from the genome-wide trends of approximately 30% in either direction (Chi-squared test, *χ*^2^ = 3529, df = 2, *P* < 2.2×10^16^; **Fig. 1d**, **top and middle**). For chromosome 20, there was relative copy number loss for the BS sample in the short arm and relative copy number gain in the long arm (**Fig. 1e**, **top)**. Consistently, we observed similar skewness of differential signals toward the sample with a relatively higher copy number: an overrepresentation of less accessible peaks for BS on the short arm (629 out of 874 peaks) and more accessible peaks on the long arm (1933 out of 3034 peaks) (**Fig. 1e**, **top and middle**). These observations strongly suggest that CNV was the main driver of the differential signals at these regions and may thus substantially confound the interpretation of the results.

ChIP-seq is another common quantitative genomic sequencing assay whose readout, like ATAC-seq, is based on quantifying the number of reads/fragments from a genomic interval. Here, we performed G4 ChIP-seq using antibodies against a secondary DNA structure, G-quadruplexes (G4), and mapped the endogenously formed G-quadruplexes in the aforementioned WT and BS cell lines. G-quadruplexes are a four-stranded secondary DNA structure formed by G-rich sequences through cyclic Hoogsteen hydrogen bonds and are one of the DNA substrates of BLM helicase [32, 33]. In total, we identified 20,072 high-confidence G4 peaks and performed the differential analysis to characterize differential G4 forming sites via *DiffBind*. We detected 11,730 (58.5%) significantly differential sites (FDR < 0.05), of which 6217 and 5513 showed increased and decreased signals, respectively. Similar to our previous observation for ATAC-seq, we observed an even stronger CN-dependent bias in the differential signals **(Additional file 2: Fig.S1a)**. Peaks identified to have decreased G4 forming signals were overrepresented in regions with relatively lower copy numbers in BS vs. WT (log_2_ CNR <0) and vice versa.

Our results from both ATAC-seq and ChIP-seq showed that copy number differences between contrasted samples could drive the differential signals. Notably, while BS cells have been reported to have excessive sister chromatid exchange (SCE) events, tissues/cells directly sampled from BS individuals mostly showed normal karyotypes [26, 27, 34]. Occasional chromosome losses has been reported in some studies though, rarely has BS been directly linked to complex aneuploidy and broad genome-wide chromosomal aberrations in normal tissues, as observed in our cell lines [26, 34–36]. Moreover, the fibroblast cell line derived from a healthy donor showed widespread chromosomal aberrations. Therefore, we interpret the chromosomal aberrations in the fibroblast cell lines to be an artifact during the immortalization of the cell line rather than a primary effect of BS. Misattributing the variation driven by copy number variation (CNV) between WT and BS samples as differential signals due to BS would obscure the biological significance of the observed differences.

### Copy number normalization separates differential signals driven by copy number variation

We have demonstrated the impacts of copy number variation on differential analysis. If left uncorrected, CNV could mask biologically relevant differential signals and confound the interpretation of the results. It is therefore necessary to separate the differential peaks driven by CNV and/or aneuploidy from other factors, particularly, the factor of interest, here BS. This is also of general interest because aneuploidy and CNV are common to various biological materials, e.g., cancer tissues. To account for copy number in the differential analysis, we implemented CN normalization after signal quantification, but before data normalization in the pipeline (**Fig. 2**).

**Fig. 2.**
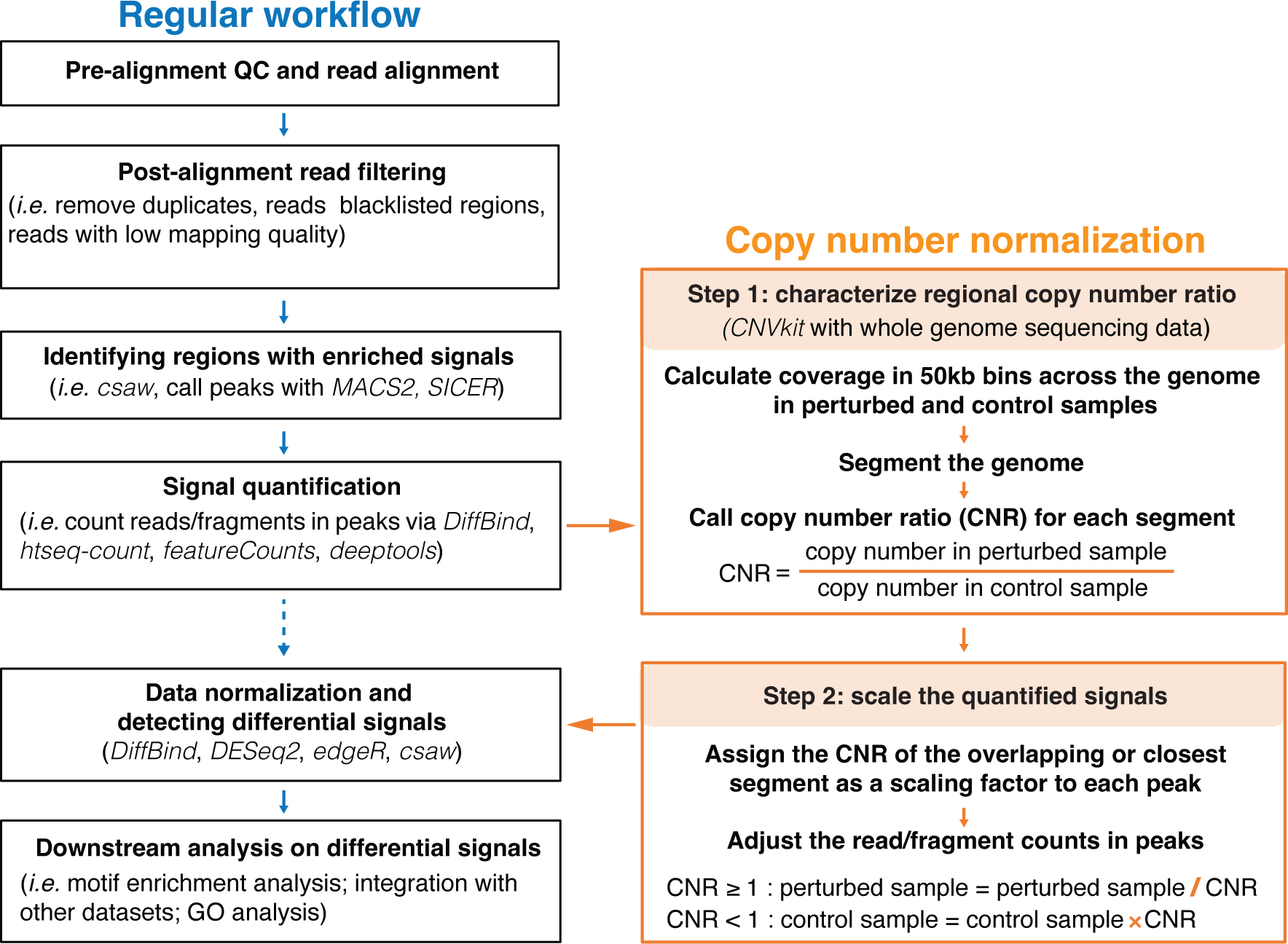
A copy-number-aware differential analysis pipeline featuring copy number normalization.

Copy number normalization mainly consists of two steps, estimating and then correcting differences in copy number. The first step is to characterize the copy number ratio, which is the regional relative copy number variation in the perturbed sample relative to the control sample. Notably, we aim to detect CNR instead of an absolute genome-wide CN determination within each sample, because it does not require prior knowledge of the karyotype in any of the contrasted samples. Here, we used *CNVkit* to estimate local CNR [30]. It uses genomic sequencing data from ATAC-seq (or ChIP-seq input) from both BS and WT samples to first calculate the coverage in non-overlapping 50kb bins in the genome, then segment the genome, and lastly assess the CNR for each segment. Next, we assigned each peak to its overlapping DNA segment or the closest DNA segment and corrected the the read/fragment count in this peak by applying the CNR of the DNA segment as a scaling factor. The modified count matrix is used in the subsequent steps of data normalization and calling of differential signals (**Fig. 2**). Specifically, for peaks in regions with copy number gain (log_2_ CNR > 0), the read/fragment counts in BS samples were divided by CNR whereas for peaks in regions with copy number loss (log_2_ CNR < 0), the read/fragment counts in WT samples were multiplied by CNR to avoid inflating the statistical power to detect the differential signals. Notably, these add-on steps can be easily integrated into most pipelines using a count-based approach for signal quantification.

After having applied CN normalization, we re-analyzed the ATAC-seq data between BS and WT samples and obtained strikingly different results **(Fig. 3a, left)**. Importantly, the copy number-dependent bias of differential signals was largely removed. This effectively eliminated the overrepresentation of differential signals towards the sample with a relatively higher copy number in the region (**Fig. 1c**, **right**; **Fig. 1e**, **bottom**). For instance, on chromosome 17 with relative copy number gain, fewer peaks remained significantly differential following CN normalization (3559 vs. 5238 without CN normalization) and differential peaks displayed a more balanced distribution (1822 increased and 1737 decreased) along the chromosome **(Fig. 1d**, **bottom**; **Fig. 3a, middle)**. Similarly, on the long arm of chromosome 20, the overrepresentation of increased signals is attenuated (25.18% vs. 63.7% without CN normalization) **(Fig. 3a, right)**. Out of 1348 peaks on the short arm with copy number loss, the significantly differential signals are comparable before and after CN normalization (775 vs. 825) (**Fig. 1e**, **bottom**; **Fig. 3a, right**). We also noted that on the short arm, a large majority of the significantly less accessible regions retained their differential status after CN normalization, suggestive of intrinsic regulatory changes in these regions in BS. Alternatively, this could also be attributed to relatively small copy number differences in these regions, resulting in less CNV-driving bias in the differential signals.

**Fig. 3.**
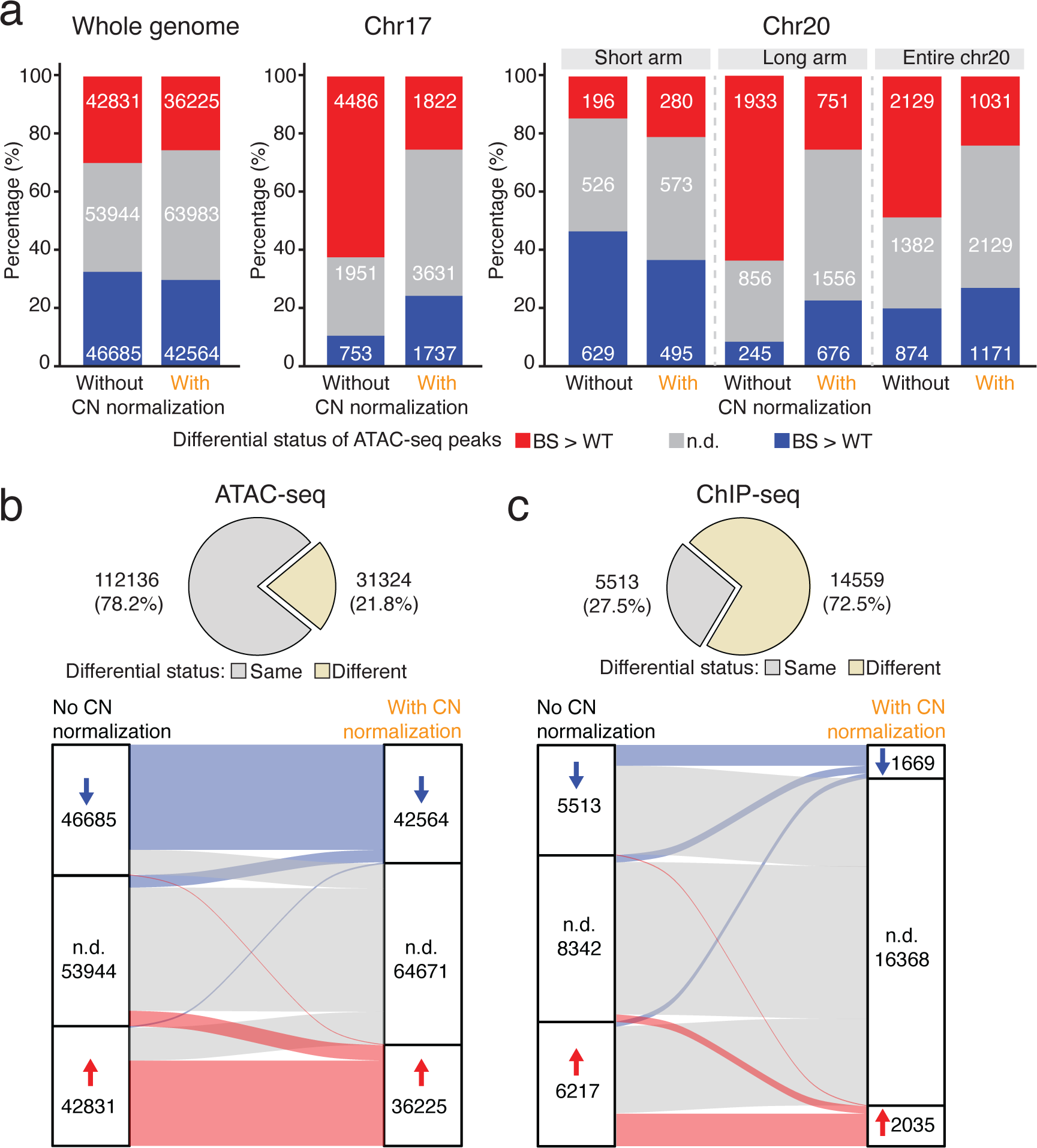
Impacts of copy number normalization on differential analysis. **a** Number of significantly and not significantly differential ATAC-seq peaks across the genome (left), on chromosome 17 (chr17, middle) and on chr20 (right) before and after applying copy number (CN) normalization. N.d. indicates not significantly differential signals with adjusted *P* (p-adj) ≥ 0.05. **b** Differential status of open chromatin regions before and after applying CN normalization. N.d. indicates not significantly differential signals with p-adj ≥ 0.05. **c** Differential status of G-quadruplex forming sites before and after applying CN normalization in ChIP-seq data. N.d. indicates not significantly differential signals with FDR ≥ 0.05

To characterize the global impact of CN normalization on the outcome of differential analysis, we determined how the differential status has changed for each open chromatin region in ATAC-seq data or G4 peaks in ChIP-seq data following CN normalization. Overall, about 20% of the ATAC-seq peaks changed their differential status, with most of these now re-classified as non-differential (adjusted *P* ≥ 0.05; **Fig. 3a, left**; **Fig. 3b**). Notably, the CN normalization has remarkably stronger impacts on the ChIP-seq data. The number of significantly differential G4 forming sites dropped by 68% to only 3704 from 11730 without CN normalization (**Fig. 3c**). These observations suggest that differential signals identified with the usual workflow are substantially confounded by copy number variation, particularly in ChIP-seq data. Crucially, we noted that our ChIP-seq data had a lower signal-to-noise ratio compared to the ATAC-seq data (**Additional file 2: Fig. S1b**). This has implications for signal quantification. This is because during counting reads/fragments at peaks, the increased background signal could also be counted as signals. Importantly, this contribution from background reads increased in proportion to the copy number. As a result, data with worse signal-to-noise ratios would be biased to a larger extent by the differences in copy numbers between samples. Consistent with this effect, peaks in regions with extreme CNR were prone to be identified as differential upon no CN normalization.

### Copy number normalization identifies accessible chromatin regions with dosage effects and compensatory effects in Down syndrome

So far, we have examined a case where CN differences is not thought to be the primary disease-causing mechanism. Next, we investigated the impact of CN differences in Down syndrome (DS), a classical example of aneuploidy. Down syndrome, also known as trisomy 21, is a genetic disorder caused by harboring an extra copy (95% of the cases) of chromosome 21. It is the most common chromosomal anomaly and the most common genetic cause of significant intellectual disability [37, 38]. We used publicly available ATAC-seq data from paired fraternal lymphoblasts cell lines from DS (47, XY, +21) and non-DS (46, XY) individuals (referred to as wildtype or WT in the subsequent text) [39].

In total, we identified 1397 open chromatin regions on chromosome 21. No open chromatin region was in the 34 kb highly restricted Down syndrome critical region (hg38, chr21: 37,929,229 - 37,963,130) (**Additional file 2: Fig. S3**), the minimal region whose triplication is shared by all DS subjects and is absent in all non-DS subjects [40]. In general, open chromatin regions on chromosome 21 displayed higher chromatin accessibility in the DS sample compared to the euploid WT sample. By contrast, on chromosome 17, where both DS and WT samples are diploid, we observed comparable signals between WT and DS samples (**Fig. 4a**, **4b**). This observation is consistent with our observation with BS vs. WT samples, that elevated chromatin accessibilities were associated with a relatively higher copy number.

**Fig. 4.**
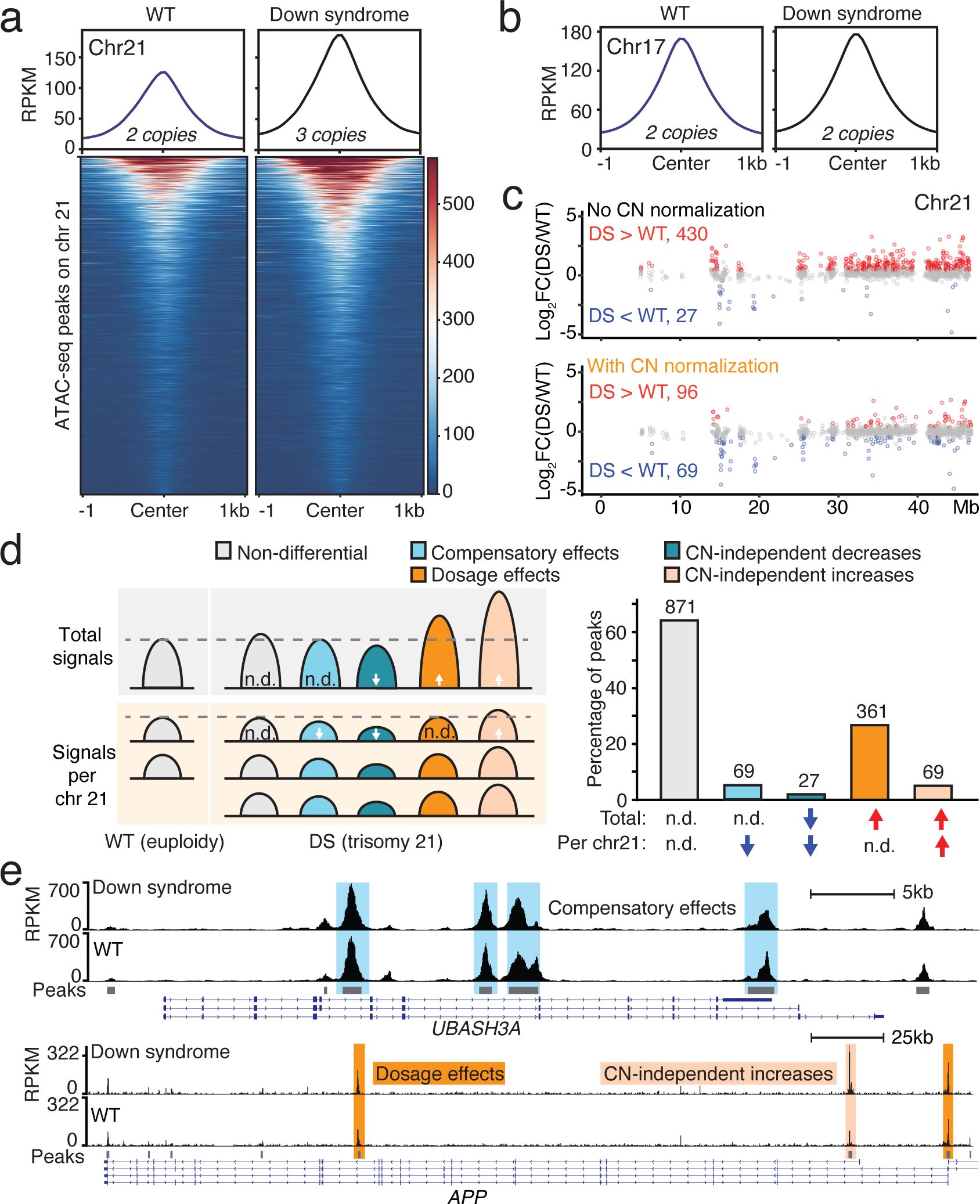
Copy number normalization identifies regions with dosage and compensatory effects in Down syndrome. **a** Average ATAC-seq signal profiles (top) and chromatin accessibility (bottom) of peaks on chromosome 21 (chr21) in wildtype (WT) and Down syndrome (DS) samples. **b** Average ATAC-seq signal profiles of peaks on chr17 in WT and DS samples. **c** Distribution of differential signals regarding total chromatin accessibility (top) and average chromatin accessibility per chr21 (bottom) in ATAC-seq peaks. **d** Categories of open chromatin regions on chr21 defined according to changes in total chromatin accessibility and average chromatin accessibility per chr21. N.d. indicates not significantly differential signals with adjusted *P* (p-adj) ≥ 0.05; arrows denote significantly increased or decreased signals with p-adj < 0.05. **e** Open chromatin regions in *UBASH3A* and *APP* genes exhibiting compensatory effects (shaded in blue), dosage effects (shaded in dark orange) or both dosage effects and CN-independent increases (shaded in light orange). Peaks without shading displayed no change in either total or average chromatin accessibility per chr21.

Here, aneuploidy is the central cause of DS: the differential signals due to CNV reflect the direct impact of copy number gain of chromosome 21 and are thus biologically relevant. Therefore, we performed the genome-wide differential analyses with and without CN normalization. They produced strikingly different results for peaks on chromosome 21. Without applying CN normalization, trisomy 21 overwhelmingly drives the differential accessibility signal on chromosome 21. Out of the 457 significantly differential accessible regions, 430 (94.1%) regions showed increased chromatin accessibility while only 27 regions with decreased accessibility (**Fig. 4c**, **top)**. With CN normalization, we identified only 165 significantly differential signals with 69 displaying increased signals and 96 showing decreased signals, including the same 27 regions that exhibited decreased signals without CN normalization. (**Fig. 4c**, **bottom)**.

In the context of DS, differential signals identified by these two approaches call for distinct biological interpretations. One direct impact of trisomy 21 is the gene dosage and potential dosage effects which are determined by the amount of expressed gene product. There have been extensive studies in DS reporting increased transcription and protein expression levels of genes on chromosome 21, demonstrating gene dosage effects [41–46]. Meanwhile, in many other biological contexts, mechanisms engage to prevent expression levels from increasing linearly with CN, e.g., dosage compensation [41, 45, 47, 48]. Here, we should look beyond expressed gene product, and extend these concepts to regulatory signals of chromatin such as chromatin accessibility and chromatin binding. Using ATAC-seq data and chromosome 21 in DS as an example, the standard differential analysis pipeline compares the total signals from three gene copies in the DS sample with those from two gene copies in the WT sample (**Fig. 4d**, **left**). This approach reveals differential signals linked to the presence of an extra chromosome 21. For instance, regions with increased signals imply dosage effects of chromatin accessibility. In contrast, within the CN-aware pipeline incorporating CN normalization, the average signal per-chromosome copy is compared. Specifically, the total signal from all three copies in DS samples is divided by the copy number ratio (CNR = 3/2). Subsequently, the aggregated chromatin accessibility adjusted for two gene copies is compared against the signal from WT to reveal CN-independent changes (**Fig. 4d**, **left**). For instance, regions with decreased signals suggest dosage compensation of chromatin accessibility in response to trisomy 21.

As discussed above, these two approaches identify CN-dependent and -independent differential signals in DS. Based on the combination of the changes in total and per-chromosome copy signals, peaks on chromosome 21 could be divided into the following categories: Ⅰ) regions with no change; Ⅱ) regions with over-compensatory effects and CN-independent decreases, showing decreased total and per-copy signal; Ⅲ) regions with compensatory effects, in which total signals remain unchanged but per-copy signal decreases; Ⅳ) regions showing only dosage effects, with increased total signals but no change in per-copy signal; Ⅴ) regions with both dosage effects and CN-independent increases (**Fig. 4d**, **right**). Out of 940 regions showing no change in total chromatin accessibility in DS (category Ⅰ and Ⅱ), the conventional pipeline would misidentify all of them as regions with compensatory effects, whereas CN normalization enabled us to show that only 69 (7.3%) have compensatory effects (**Fig. 4d**, **right**). Moreover, among all the regions with increased total chromatin accessibility in DS (category Ⅳ and Ⅴ), CN normalization distinguished that approximately 80% are purely due to dosage effects, with the rest exhibiting further CN-independent increases (**Fig. 4d**, **right**).

Many of the identified differential signals are in genes associated with the clinical symptoms of DS. It has been reported that DS individuals are more susceptible to Alzheimer’s disease as the gene encoding amyloid precursor protein, *APP*, is located on chromosome 21 [37, 38]. The peak in the promoter (in this study defined as TSS ± 3kb) of *APP* showed only dosage-dependent increases whereas another peak in the promoter of a splicing variant showed a further increase in the per-copy signal, which may further boost the expression of *APP* and that specific transcript variant (**Fig. 4e**, **top**). Additionally, DS individuals exhibit higher risks of autoimmune diseases and developing severe responses to infectious diseases [37, 38, 49–51]. Tissues and cells from DS individuals and mouse models suggest that DS exhibits interferon hyperactivity and chronic inflammation [42, 52, 53]. One of the major reasons for dysregulated immune function in DS is that chromosome 21 carries genes essential for immune function. A T-cell ubiquitin ligand coding gene on chromosome 21, SH3 domain-containing protein A (*UBASH3A*), negatively regulates T-cell signaling and it is a risk gene for autoimmune diseases [54]. Almost all the open chromatin regions in this gene displayed a decreased average signal on each chromosome 21 such that the total chromatin accessibility remained the same in DS (**Fig. 4e**, **bottom**). Moreover, four of the six interferon receptor genes are also on chromosome 21. Among the 24 peaks in this interferon receptor gene cluster, 14 showed increased chromatin accessibility due to dosage effects, and one peak in the intronic regions of *IFNAR1* and two peaks in the intergenic regions showed CN-independent changes (**Additional file 2: Fig. S4a**). Another gene on chromosome 21 that is associated with inflammatory signals in the brain in DS is *S100B* gene [38]. Despite our data generated from lymphocytes rather than from brain tissues, ATAC-seq peaks near this gene showed dosage effects and increased chromatin accessibility per copy (**Additional file 2: Fig. S4b**). The above examples illustrate how combining overall chromatin accessibility with per-allele regulatory changes can further help explain DS phenotypes apart from transcription dosage effects. These CN-independent regulatory changes would have been missed without incorporating the CN normalization step.

As further evidence on the robustness of the procedure, we noted that the vast majority of differential signals on the other chromosomes except chromosome 21 remained the same (**Additional file 2: Fig. S2b, 2c and 2d**), suggesting that the current pipeline is specific in its ability to correct for effect due solely to local copy number differences between contrasted samples.

## Discussion

Here we have demonstrated that copy number variation among contrasted samples, a factor often overlooked in the differential analysis, can and does drive much of the observed differential signal. Given the prevalence of aneuploidy in diseases such as cancer and cell lines used in biomedical research, such effects could hold clinical relevance. To address this, we developed a CN-aware pipeline featuring copy number (CN) normalization, which accounts for the copy number differences between samples and adjusts the signals to the same copy number before downstream data normalization. We presented two cases in biomedical studies to illustrate the impacts of copy number differences on the differential analysis. We further demonstrated that applying our proposed CN-aware pipeline can effectively segregate the CN-dependent from the CN-independent differential signals.

In the case of Bloom syndrome, caused by loss-of-function mutations in *BLM*, cell lines derived from both the healthy donor and the BS individual accumulated complex chromosomal aberrations irrelevant to the disease status. We observed that without accounting for CNV between the samples, differential signals are biased toward the sample with a higher copy number. The copy number differences affect approximately 20% of differential signals in ATAC-seq and 70% in ChIP-seq data. These false-positive signals could be attributed to BS, leading to potential misinterpretation of results. By applying CN normalization, we largely removed the CN-dependent differential signals. In the case of Down syndrome, trisomy 21 is the central cause of the disorder and thus the main factor of interest. But by performing differential analyses with and without CN normalization, we were able to detect CN-dependent and -independent changes. Combining results from both approaches enables a nuanced biological interpretation of the changes in chromatin accessibility. By extending the concepts of dosage effects and dosage compensation to chromatin accessibility, we revealed the subset of peaks with genuine dosage compensatory effects and distinguished peaks with only dosage effects from those further CN-dependent increases.

Such regions with CN-independent changes would have been overlooked without considering CNV, yet they are potentially clinically relevant. For instance, most of the peaks in the risk gene for Alzheimer’s disease showed increased chromatin accessibility mainly due to the extra copy of chromosome 21. One peak in the promoter regions of a transcript variant of *APP* gene exhibited further CN-independent increases, potentially boosting the expression of a specific variant of the APP protein and contributing to increased risk of Alzheimer’s disease in individuals with DS. Additionally, most of the peaks in the overexpressed interferon receptor genes showed pure dosage effects [42, 49, 52]. Conversely, in an autoimmune-related gene, *UBASH3A*, nearly all the peaks in this gene showed decreased chromatin accessibility, possibly in compensation for the third gene copy, implying a critical role and tight regulation of this gene. These CN-independent changes plausibly arise secondarily from the primary molecular changes upon trisomy 21 and provide novel insights into dosage imbalance and the pathogenesis of DS. Further studies on these regions are required to understand the disrupted epigenetic regulatory network. In particular, performing single-cell assays on DS samples to tangle the activation and silencing of different chromosome 21 copies will provide deeper insights into dosage and compensatory effects of chromatin accessibility and gene products, which help understand the changes in regulatory network in DS.

Notably, instead of determining the exact CNV within each sample, our CN normalization aims to characterize and utilize the regional copy number ratio between the contrasted samples. Detecting the former can be challenging as it requires prior knowledge of the karyotypes, which may not be readily available, especially for samples with complex CNV and aneuploidy, such as cancer cells. In contrast, characterizing the CNR circumvents this issue, although there are scenarios where direct per-sample CNV determination may be preferred. Here we used *CNVkit* with a bin size of 50kb to estimate CNR [30]. This tool estimates the copy number by comparing the coverage of samples to a reference sample in each bin and then segments the genome based on variation in the coverage. The resolution of CNR can be improved by deeper sequencing and using smaller bin sizes. Alternatively, *CNVkit* can also call the exact CNV within each individual sample. Additionally, *CNVkit* offers a convenient all-in-one command. Nevertheless, other tools such as *CopywriteR*, *QDNAseq* and pipelines like the Genome Analysis Toolkit (*GATK*) best practice for Somatic copy number variant discovery can also be employed to characterize CNV and calculate CNR [55– 59]. Once CNR is obtained, assigning CNR as scaling factors to peaks and adjusting the quantified signals do not rely on any specific bioinformatic tools. These steps are not exclusive to our workflow or our choices of bioinformatic tools; instead, they can be implemented independently as add-on steps to most differential analysis pipelines for ATAC-seq and ChIP-seq.

Our pipeline mainly focuses on addressing the impacts of CNV in differential analysis and not in the preceding step of identifying regions with enriched signals or peak calling. Incorporating tools like Histone Modification in Cancer (*HMCan*), which uses Hidden Markov Model (HMM) and accounts for copy number variation to call peaks, could be integrated into our proposed pipeline to eliminate false peaks and further improve the results [60]. The same group also developed *HMCan-diff* for differential analysis counting for copy number variation [61]. While we agree with taking CNV into account, this tool differs from our recommendation in that it aims to be a stand-alone pipeline. As such, this tool may limit the flexibility of using other statistical models in data normalization methods and testing for differential signals. Apart from ATAC-seq and ChIP-seq, variation in copy number, being the baseline of DNA substrate, will also be a significant factor in the differential analysis of other functional genomic assays that employ count-based approaches to quantify signals from DNA and/or chromatin, like DNase I hypersensitive sites sequencing (DNase-seq), micrococcal nuclease digestion with deep sequencing (MNase-seq), Cleavage Under Targets and Release Using Nuclease (CUT&RUN), Cleavage Under Targets and Tagmentation or (CUT&Tag). Notably, our CN normalization strategy is not limited to ATAC- and ChIP-seq but can also be applied to the aforementioned data types. In our implementation, the CN normalization steps can be seamlessly integrated into the differential analysis pipelines of these assays as add-on steps.

## Conclusions

We demonstrated that CNV between contrasted samples is a significant factor in the differential analysis of count-based functional genomic assays, such as ATAC-seq or ChIP-seq. Copy number differences could drive the differential signals. To address this, we proposed a CN-aware differential analysis pipeline featuring CN normalization. The underlying core concept for CN normalization is that the signal from a specific genomic region is the aggregated signal from all gene copies. In the case of BS where CNV is an artifact rather than a biologically relevant factor, incorporating CN normalization into a conventional CN-unaware pipeline effectively eliminated the majority of differential signals driven by CNV, thus preventing misidentifying them as disease-related alterations. Conversely, in the case of DS with CNV being a central factor of interest, performing differential analysis with and without copy number normalization reveals signals with dosage effects and compensatory effects, providing a new perspective to interpret the results. Importantly, both the concept and the CN normalization approach apply to other data types like DNase-seq, MNase-seq, CUT&RUN, and CUT&Tag. In summary, we would like to highlight the impacts of CNV on the differential analysis of functional genomic assays, along with its underlying biological interpretation, and emphasize that CN normalization should constitute a key step in the differential analysis in general, even in studies where CNV and aneuploidy may not be central factors of interest.

## Methods

### Cell culture

Sex-matched and roughly age-matched fibroblast cell lines from Bloom syndrome and healthy donors were obtained from Coriell Institute (BS, GM08505; WT, GM00637). Fibroblast lines were cultured in 1× DMEM (Thermo Scientific, Gibco, Cat. #11960) supplemented with 1% minimum essential medium non-essential amino acids (MEM NEAA; Thermo Scientific, Gibco, Cat. #11140) and 1% Penicillin-Streptomycin (Thermo Scientific, Gibco, Cat. #15140122) and 10% or 15% fetal bovine serum (FBS; Thermo Scientific, Gibco, Cat. #10500064), respectively. Cell passaging was performed according to recommendations by Coriell Institute. All cells were cultured at 37°C under 5% CO_2_.

To collect cells, fibroblasts were first trypsinized (Trypsin-EDTA solution, Sigma-Aldrich, Cat. #T4049), pelleted at 300 g for 5 min and washed once with cold 1× Dulbecco’s phosphate-buffered saline (PBS, Thermo Scientific, Gibco, Cat. #14190).

### ATAC-seq library preparation

Cells were collected as described above and counted (automated cell counter Countess 3, Thermo Scientific; Countess™ Cell Counting Chamber Slides, Thermo Scientific, Cat. #C10283). For each reaction, 50,000 cells were subjected to omni-ATAC-seq preparation as described previously with a modified tagmentation reaction [62]. Nuclei were resuspended in 50 μl of tagmentation mix consisting of 10 μl 5× TAPS-DMF buffer (50 mM TAPS, 25 mM MgCl_2_, 50% (v/v) DMF), 3 μl Tn5 transposase [63], 16.5 μl 1x PBS, 0.25 μl 2% Digitonin, 0.5 μl 10% (v/v) Tween 20 and 19.75 μl H_2_O. DNA purified from the transposed product was then amplified via PCR (10 cycles) using Q5 High-Fidelity DNA Polymerase (New England Biolabs, Cat. #M0491L).

### G-quadruplex ChIP-seq protocol and library preparation

#### Chromatin preparation

Chromatin preparation was performed essentially as previously described with a few modifications [29]. Cells from two full 15cm dishes were initially rinsed once in 1× PBS and subsequently fixed in the dish with 1% formaldehyde in cell culture medium at room temperature for 8.5 minutes. Nuclei were then lysed in 25 μl lysis buffer and sheared in small tubes (microTUBE AFA Fiber Pre-Slit Snap-cap 6×16mm, Covaris, Cat. #520045) using an S220 instrument (Covaris). The shearing parameters were as follows: *water level = 15*, *duty cycle = 15*, *intensity = 6*, and *cycle per burst = 200*. Each chromatin sample underwent shearing for a total of 3 minutes, with a 30-second pause for every one minute of shearing.

#### G-quadruplex ChIP-seq

For each sample, 2 biological replicates, each of which with 3 technical replicates of immune-precipitation (IP) were prepared. The IP reaction was performed as described previously [29], with the following modifications: during IP with the antibody called BG4 against DNA G-quadruplexes structures, chromatin was incubated with the antibody called BG4 against DNA G-quadruplexes structures for an extended duration of 2 hours, and 10ul of Anti-FLAG M2 Magnetic Beads (Sigma-Aldrich, Cat. #M8823) was used for each reaction in pull-down. To purify DNA from both the IP samples (DNA pulled down by the beads) and the input samples (sheared chromatin without going through IP steps) after 5 times of washing the beads with WASH buffer (100 mM KCl, 0.1% (v/v) Tween 20, 10 mM Tris, pH 7.4) to remove residual nonspecific bound chromatin, 75 μl of reverse-crosslinking buffer (0.2% SDS, 1× TE, and 50 mM NaCl) was introduced. The samples were incubated at 37°C for one hour, followed by an overnight incubation at 65°C. After an additional hour of proteinase K digestion at 65°C, DNA purification was carried out using a MinElute kit (QIAGEN, Cat. #28006). The subsequent steps for library preparation were carried out with the DNA Thruplex kit (Takara, Cat. #R400674, Cat. #R400665) following the manufacturer’s protocols.

#### G-quadruplex ChIP-qPCR and library preparation

Prior to library preparation, qPCR was performed to evaluate G-quadruplex enrichment in IP vs. input (DNA extracted from sonicated chromatin without IP). We used 2X CFX SYBR Mix (Applied Biosystems, Cat. #4472942) and a set of primers targeting known G4 positive and negative control sites [29].

### Sequencing and data analysis

One biological replicate of G4 ChIP-seq libraries was sequenced on a HiSeq3000 platform (150 bp paired-end sequencing; Illumina Inc.; the Genome Core Facility at the Max Planck Institute for Biology, Tübingen). ATAC-seq libraries and the second biological replicate of G4 ChIP-seq libraries were sequenced on a NovaSeq6000 platform (150 bp paired-end sequencing; Illumina Inc.; service provider: GENEWIZ, Leipzig).

A more detailed step-by-step description and pipeline can be found both in supplementary files and on GitHub, https://github.com/Dingersrun/Copy-number-normalization

#### Read alignment and read filtering

Raw FASTQ reads were extracted with *bcl2fastq* (version 2.20). Tn5 and TruSeq adapter sequences, G repeats due to two-color base calling errors and reads shorter than 20 bp were trimmed and removed using *cutadapt* (version 4.0) [64]. Reads were then aligned to the human reference genome version hg38 using *bwa mem* (version: 0.7.17-r1188). Duplicates were marked and removed using *Picard* (https://broadinstitute.github.io/picard/, version 2.18.25). Prior to calling peaks, reads mapped to mitochondrial, reads with a mapping quality lower than 20 and reads in hg38 blacklisted regions (downloaded from http://mitra.stanford.edu/kundaje/akundaje/release/blacklists/hg38human/hg38.blacklist.be <u>d.gz</u>) were discarded.

#### Peak calling and differential analysis for ATAC-seq

ATAC-seq peak calling was performed using *MACS2* (version 2.1.1.20160309) with *callpeak –format BAMPE --nomodel –min-length 100 narrowPeak* parameters using a bamfile made from pooling equal number reads from each sample [10, 65]. The number of fragments in each ATAC-seq peak was counted with *htseq-count* (*HTSeq* version 0.9.1) [15]. Subsequently, the count matrix was normalized and differential analysis was carried out in *DESeq2* (version 1.30.1) [31]. In the case of applying copy number normalization, prior to data normalization the count matrix was adjusted as described in the following session.

#### Copy number normalization

A detailed step-by-step description can be found in the additional file 3.

The first step of copy number normalization is characterizing CNR. Genomic sequencing data from cell lines GM08505 (BS) and GM00637(WT) were used in *CNVkit* following its recommended copy number calling pipeline *(--method wgs --target-avg-size 50000*) to identify copy number alteration in BS relative to WT [30]. Gapped regions in the human genome assembly marked by Ns (http://hgdownload.soe.ucsc.edu/goldenPath/hg38/database/gap.txt.gz) were excluded from the analysis. In the output *CNVkit*, log_2_-transformed values representing CNR for genomic segments were retrieved and converted back to the original value. The second step is assigning each peak to its overlapping DNA segment or the closest DNA segment. The CNR of this segment was then used as a scaling factor to modify the read/fragment count in this peak. Specifically, for the BS vs. WT comparison, if the CNR is greater than or equal to 1 (CNR ≥ 1), the fragment counts in peaks in BS were divided by the CNR. Conversely, if the CNR was smaller than 1 (CNR < 1), the fragment counts in peaks in WT were multiplied by the CNR. For DS vs. WT comparison, the fragment counts in peaks for peaks on chromosome 21 were simply divided by 1.5 in the DS sample.

#### Peak calling and differential analysis for ChIP-seq

For each biological replicate, a pooled file was generated by subsampling 35 million reads from each technical replicate. The pooled file was subjected to peak calling using *MACS2* (version 2.1.1.20160309) with *callpeak – format BAM narrowPeak* [65]. Peak sets from 2 biological replicates were ranked by their signal values and then filtered based on reproducibility using *IDR* (version 2.0.3) [66]. The final high confidence peaks for each sample were obtained with the criteria of irreproducible discovery rate < 0.05. Peaks were also called with input samples using the same parameters, which were later used as regions to be excluded in the differential analysis in *DiffBind* (version 3.0.15) [18, 19]. The differential analysis was carried out by using default parameters with *summits=FALSE, bFullLibSize=FALSE* and *edgeR* normalization [67]. In the case of applying copy number normalization, the read counts in peaks for each sample were extracted from *DiffBind* and modified in the same way as it was for ATAC-seq. A more detailed step-by-step description can be found in supplementary files.

## Supplementary Information

Additional file 1. Copy number ratio in Bloom syndrome cell line relative to wild type cell line for each chromosome.

Additional file 2. Supplementary figures.

Additional file 3. Step-by-step pipelines for differential analysis with and without copy number normalization for ATAC-seq and ChIP-seq data.

## Supporting information

Supplemental Data 1 - CNR per chromosme

Supplementary Data 2 - DifferentialAnalysis CNV pipeline

## Acknowledgments

We thank Detlef Weigel and Marja Timmermans for inspiring this idea. We thank members of the Chan and Jones lab and Yinan Wang for helpful discussions and critical reading of the manuscript. We also thank the Genome Center at the Max Planck Institute for Biology Tübingen for providing support. We thank Andre Noll for computing support.

## Author contributions

D.S. conceived the idea, designed the experiment, generated the data, conducted the data analysis and developed the pipeline. Y.F.C. supervised the data collection and helped interpret the results. M.P., V.S. and Y.F.C. provided experimental or computational support. D.S. drafted the manuscript with input from all authors. All authors reviewed and approved the final version of the manuscript.

## Availability of data and materials

ATAC-seq and ChIP-seq data from WT and BS fibroblast cell lines are available at the NCBI GEO repository under accession numbers GSE259257 and GSE259258.

ATAC-seq data from Down syndrome and WT are ATAC-seq data obtained from samples cultivated at 37 °C in the study from Cardiello *et al.* under GEO accession number GSE173536 with the following entries: GSM5269423, GSM5269425, GSM5269427, GSM5269428, GSM5269429, GSM5269430, GSM5269433 and GSM5269434 [39].

## Declarations

### Ethics approval

Not applicable.

---

**Supplementary Fig. 1.**
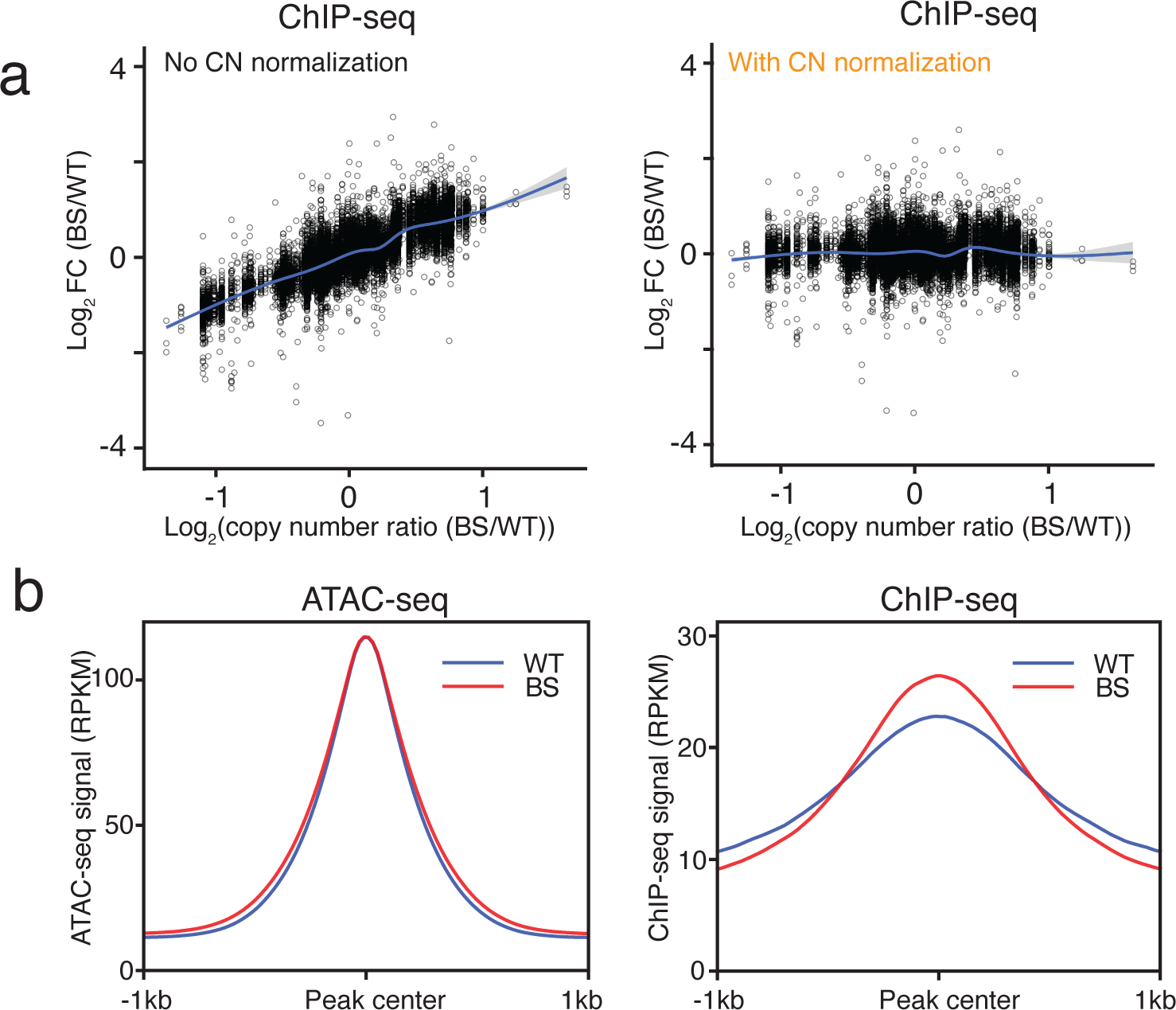
Impacts of copy number normalization on the differential analysis of ATAC-seq and ChIP-seq. **a** The trended biases of differential G-quadruplex formation from copy number differences without (left) and with (right) applying copy number normalization in ChIP-seq data. Log_2_CNR > 0 and log_2_CNR < 0 indicate relative number gain and loss in BS, respectively. The lines represent the LOESS smoothing curve for the data with the grey area indicating the 95% confidence interval for the fitted curve. **b** Average signal profiles of ATAC-seq and ChIP-seq in their corresponding peak sets.

**Supplementary Fig. 2.**
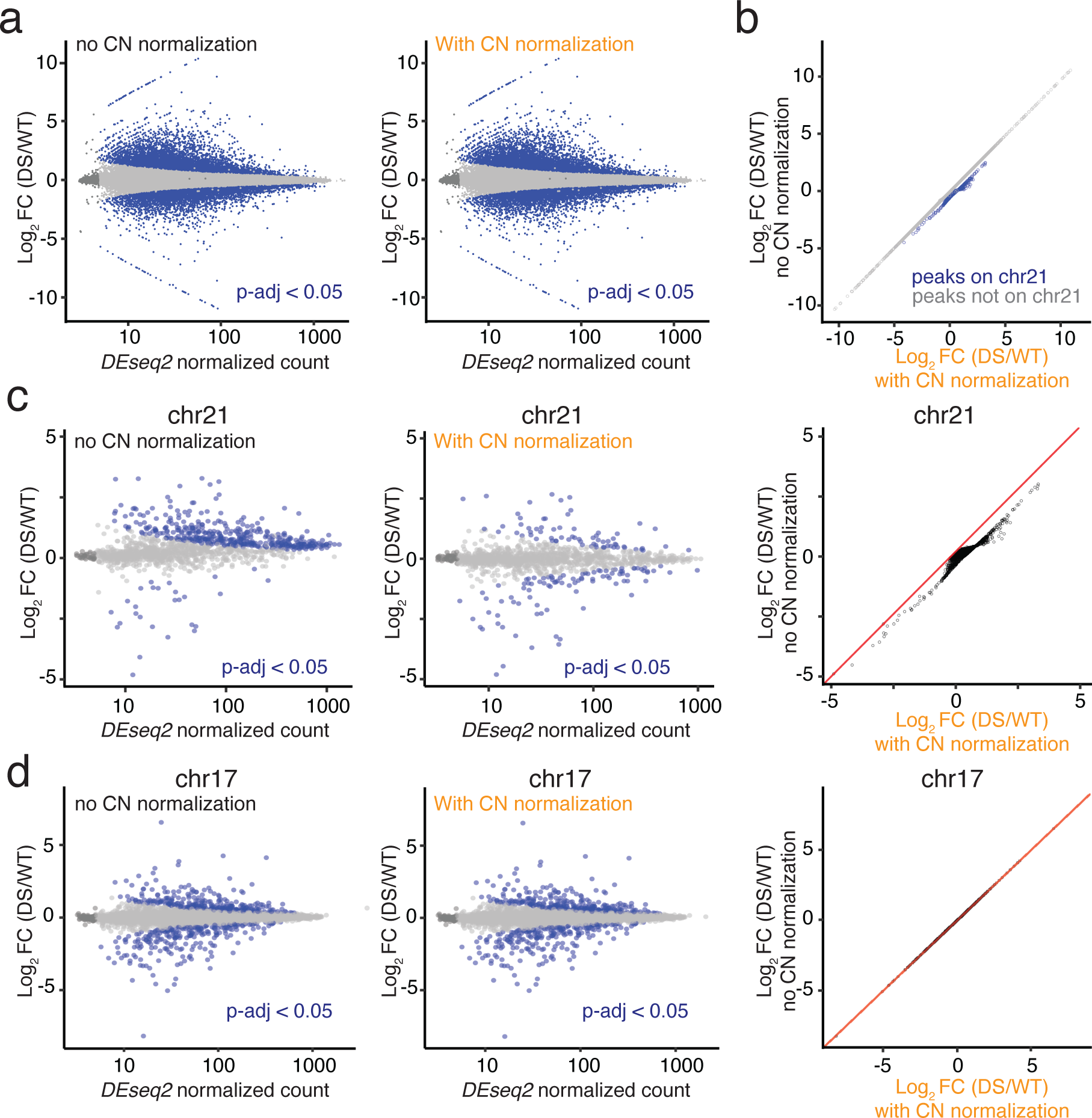
Impacts of copy number normalization on differential anlaysis of Down syndrome ATAC-seq data. **a** MA plots of the genome-wide differential signals without (left) and with (right) applying copy number (CN) normalization. Differential peaks with adjusted *P* (p-adj) < 0.05 are depicted in blue while those with p-adj ≥ 0.05 are shown in grey. **b** A comparison of the differential signals without and with applying CN normalization for peaks on chromosome 21 (chr21) (blue) and other chromosomes (grey). **c** Differential signals for peaks on chr21 without (left) and with (right) CN normalization. **d** Differential signals for peaks on chromosome 17 (chr17) without (left) and with (right) CN normalization.

**Supplementary Fig. 3.**
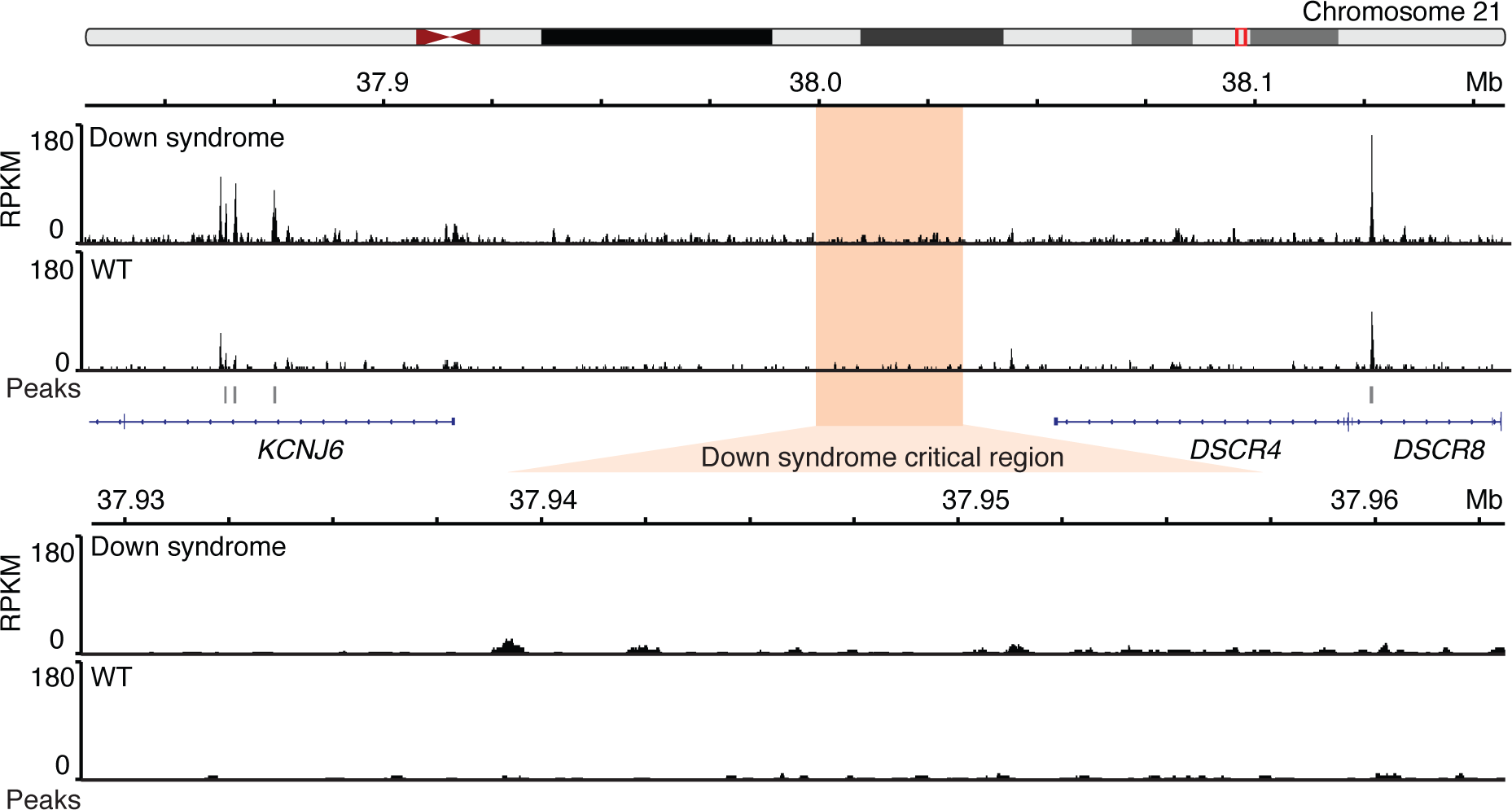
No open chromatin region in the minimal critical region in Down syndrome. The top panel is genome browser screenshot of ATAC-seq tracks in genes near the Down syndrome critical region. Bottom panel is the zoom-in view of this regions without any ATAC-seq peak.

**Supplementary Fig. 4.**
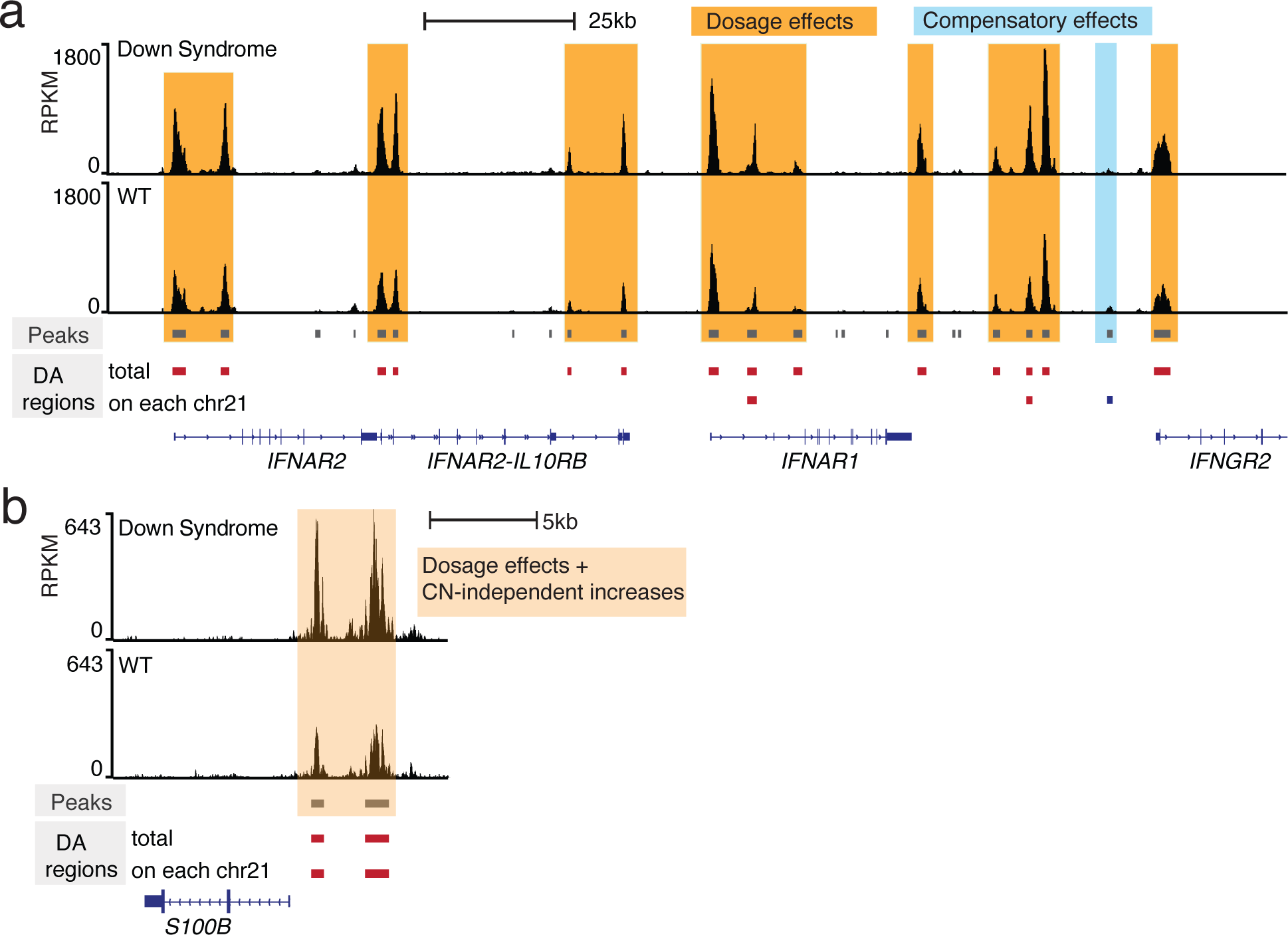
Example loci on chromosome 21 with dosage and compensatory effects in Down syndrome. **a** The genome browser screenshot of ATAC-seq tracks in interferon receptor gene clusters. Most of them showed increased total chromatin accessibility (shaded in dark orange) and one showed compensatory effects (shaded in blue). Differentially accessible (DA) regions depicted in red and blue bars represent regions with significantly increased and decreased chromatin accessibility, respectively. **b** The genome browser screenshot of ATAC-seq tracks near the transcription start site of *S100B* gene. Both showed increased chromatin accessibility in total signal and averaged signal on each chromosome 21. Red bars indicate regions with significantly increased chromatin accessibility in Down syndrome.

## References

1. Buenrostro JD, Giresi PG, Zaba LC, Chang HY, Greenleaf WJ. Transposition of native chromatin for fast and sensitive epigenomic profiling of open chromatin, DNA-binding proteins and nucleosome position. Nat Methods. 2013;10:1213–8.

2. Klemm SL, Shipony Z, Greenleaf WJ. Chromatin accessibility and the regulatory epigenome. Nat Rev Genet. 2019;20:207–20.

3. Ma S, Zhang Y. Profiling chromatin regulatory landscape: insights into the development of ChIP-seq and ATAC-seq. Mol Biomed. 2020;1:9.

4. Furey TS. ChIP–seq and beyond: new and improved methodologies to detect and characterize protein–DNA interactions. Nat Rev Genet. 2012;13:840–52.

5. Park PJ. ChIP–seq: advantages and challenges of a maturing technology. Nat Rev Genet. 2009;10:669–80.

6. Corces MR, Granja JM, Shams S, Louie BH, Seoane JA, Zhou W, et al. The chromatin accessibility landscape of primary human cancers. Science. 2018;362:eaav1898.

7. Wang J, Zibetti C, Shang P, Sripathi SR, Zhang P, Cano M, et al. ATAC-Seq analysis reveals a widespread decrease of chromatin accessibility in age-related macular degeneration. Nat Commun. 2018;9:1364.

8. Yan F, Powell DR, Curtis DJ, Wong NC. From reads to insight: a hitchhiker’s guide to ATAC-seq data analysis. Genome Biol. 2020;21:22.

9. Reske JJ, Wilson MR, Chandler RL. ATAC-seq normalization method can significantly affect differential accessibility analysis and interpretation. Epigenetics & Chromatin. 2020;13:22.

10. Gaspar JM. Improved peak-calling with MACS2. preprint. Bioinformatics; 2018.

11. Landt SG, Marinov GK, Kundaje A, Kheradpour P, Pauli F, Batzoglou S, et al. ChIP-seq guidelines and practices of the ENCODE and modENCODE consortia. Genome Res. 2012;22:1813–31.

12. Hitz BC, Lee J-W, Jolanki O, Kagda MS, Graham K, Sud P, et al. The ENCODE Uniform Analysis Pipelines. 2023;:2023.04.04.535623.

13. Quinlan AR, Hall IM. BEDTools: a flexible suite of utilities for comparing genomic features. Bioinformatics. 2010;26:841–2.

14. Ramírez F, Dündar F, Diehl S, Grüning BA, Manke T. deepTools: a flexible platform for exploring deep-sequencing data. Nucleic Acids Research. 2014;42:W187–91.

15. Anders S, Pyl PT, Huber W. HTSeq—a Python framework to work with high-throughput sequencing data. Bioinformatics. 2015;31:166–9.

16. Liao Y, Smyth GK, Shi W. featureCounts: an efficient general purpose program for assigning sequence reads to genomic features. Bioinformatics. 2014;30:923–30.

17. Lun ATL, Smyth GK. csaw: a Bioconductor package for differential binding analysis of ChIP-seq data using sliding windows. Nucleic Acids Research. 2016;44:e45–e45.

18. Stark R, Brown G. DiffBind: Differential binding analysis of ChIP-Seq peak data.

19. Ross-Innes CS, Stark R, Teschendorff AE, Holmes KA, Ali HR, Dunning MJ, et al. Differential oestrogen receptor binding is associated with clinical outcome in breast cancer. Nature. 2012;481:389–93.

20. Liang Q, Conte N, Skarnes WC, Bradley A. Extensive genomic copy number variation in embryonic stem cells. Proceedings of the National Academy of Sciences. 2008;105:17453– 6.

21. Steele CD, Abbasi A, Islam SMA, Bowes AL, Khandekar A, Haase K, et al. Signatures of copy number alterations in human cancer. Nature. 2022;606:984–91.

22. Shadeo A, Lam WL. Comprehensive copy number profiles of breast cancer cell model genomes. Breast Cancer Research. 2006;8:R9.

23. Stepanenko AA, Dmitrenko VV. HEK293 in cell biology and cancer research: phenotype, karyotype, tumorigenicity, and stress-induced genome-phenotype evolution. Gene. 2015;569:182–90.

24. Chu WK, Hickson ID. RecQ helicases: multifunctional genome caretakers. Nat Rev Cancer. 2009;9:644–54.

25. Croteau DL, Popuri V, Opresko PL, Bohr VA. Human RecQ Helicases in DNA Repair, Recombination, and Replication. Annu Rev Biochem. 2014;83:519–52.

26. Cunniff C, Bassetti JA, Ellis NA. Bloom’s Syndrome: Clinical Spectrum, Molecular Pathogenesis, and Cancer Predisposition. Mol Syndromol. 2017;8:4–23.

27. Chaganti RSK, Schonberg S, German J. A Manyfold Increase in Sister Chromatid Exchanges in Bloom’s Syndrome Lymphocytes. Proc Natl Acad Sci USA. 1974;71:4508–12.

28. Chester N, Babbe H, Pinkas J, Manning C, Leder P. Mutation of the Murine Bloom’s Syndrome Gene Produces Global Genome Destabilization. Molecular and Cellular Biology. 2006;26:6713–26.

29. Hänsel-Hertsch R, Spiegel J, Marsico G, Tannahill D, Balasubramanian S. Genome-wide mapping of endogenous G-quadruplex DNA structures by chromatin immunoprecipitation and high-throughput sequencing. Nat Protoc. 2018;13:551–64.

30. Talevich E, Shain AH, Botton T, Bastian BC. CNVkit: Genome-Wide Copy Number Detection and Visualization from Targeted DNA Sequencing. PLoS Comput Biol. 2016;12:e1004873.

31. Love MI, Huber W, Anders S. Moderated estimation of fold change and dispersion for RNA-seq data with DESeq2. Genome Biol. 2014;15:550.

32. Hänsel-Hertsch R, Di Antonio M, Balasubramanian S. DNA G-quadruplexes in the human genome: detection, functions and therapeutic potential. Nat Rev Mol Cell Biol. 2017;18:279–84.

33. Sun H, Karow JK, Hickson ID, Maizels N. The Bloom’s Syndrome Helicase Unwinds G4 DNA. Journal of Biological Chemistry. 1998;273:27587–92.

34. Chester N, Kuo F, Kozak C, O’Hara CD, Leder P. Stage-specific apoptosis, developmental delay, and embryonic lethality in mice homozygous for a targeted disruption in the murine Bloom’s syndrome gene. Genes Dev. 1998;12:3382–93.

35. van Wietmarschen N, Merzouk S, Halsema N, Spierings DCJ, Guryev V, Lansdorp PM. BLM helicase suppresses recombination at G-quadruplex motifs in transcribed genes. Nat Commun. 2018;9:271.

36. Shabtai F, Halbrecht I. Bloom’s syndrome, missing Y, hypogonadism and cancer. Clinical Genetics. 1980;18:93–5.

37. Antonarakis SE, Skotko BG, Rafii MS, Strydom A, Pape SE, Bianchi DW, et al. Down syndrome. Nat Rev Dis Primers. 2020;6:9.

38. Wilcock DM, Griffin WST. Down’s syndrome, neuroinflammation, and Alzheimer neuropathogenesis. J Neuroinflammation. 2013;10:864.

39. Cardiello JF, Westfall J, Dowell R, Allen MA. Characterizing Primary transcriptional responses to short term heat shock in paired fraternal lymphoblastoid lines with and without Down syndrome. preprint. Bioinformatics; 2023.

40. Pelleri MC, Cicchini E, Locatelli C, Vitale L, Caracausi M, Piovesan A, et al. Systematic reanalysis of partial trisomy 21 cases with or without Down syndrome suggests a small region on 21q22.13 as critical to the phenotype. Hum Mol Genet. 2016;:ddw116.

41. Hwang S, Cavaliere P, Li R, Zhu LJ, Dephoure N, Torres EM. Consequences of aneuploidy in human fibroblasts with trisomy 21. Proc Natl Acad Sci USA. 2021;118:e2014723118.

42. Waugh KA, Minter R, Baxter J, Chi C, Galbraith MD, Tuttle KD, et al. Triplication of the interferon receptor locus contributes to hallmarks of Down syndrome in a mouse model. Nat Genet. 2023;55:1034–47.

43. Liu S, Akula N, Reardon PK, Russ J, Torres E, Clasen LS, et al. Aneuploidy effects on human gene expression across three cell types. Proc Natl Acad Sci USA. 2023;120:e2218478120.

44. Stamoulis G, Garieri M, Makrythanasis P, Letourneau A, Guipponi M, Panousis N, et al. Single cell transcriptome in aneuploidies reveals mechanisms of gene dosage imbalance. Nat Commun. 2019;10:4495.

45. Antonarakis SE. Down syndrome and the complexity of genome dosage imbalance. Nat Rev Genet. 2017;18:147–63.

46. M S, C O, A B, D D, R A, K W, et al. Impact of increased APP gene dose in Down syndrome and the Dp16 mouse model. Alzheimer’s & dementia: the journal of the Alzheimer’s Association. 2022;18.

47. Conrad T, Akhtar A. Dosage compensation in Drosophila melanogaster: epigenetic fine-tuning of chromosome-wide transcription. Nat Rev Genet. 2012;13:123–34.

48. Xing Z, Li Y, Cortes-Gomez E, Jiang X, Gao S, Pao A, et al. Dissection of a Down syndrome-associated trisomy to separate the gene dosage-dependent and -independent effects of an extra chromosome. Human Molecular Genetics. 2023;32:2205–18.

49. Chung H, Green PHR, Wang TC, Kong X-F. Interferon-Driven Immune Dysregulation in Down Syndrome: A Review of the Evidence. J Inflamm Res. 2021;14:5187–200.

50. Malle L, Patel RS, Martin-Fernandez M, Stewart OJ, Philippot Q, Buta S, et al. Autoimmunity in Down’s syndrome via cytokines, CD4 T cells and CD11c+ B cells. Nature. 2023;615:305–14.

51. Illouz T, Biragyn A, Iulita MF, Flores-Aguilar L, Dierssen M, De Toma I, et al. Immune Dysregulation and the Increased Risk of Complications and Mortality Following Respiratory Tract Infections in Adults With Down Syndrome. Front Immunol. 2021;12:621440.

52. Sullivan KD, Lewis HC, Hill AA, Pandey A, Jackson LP, Cabral JM, et al. Trisomy 21 consistently activates the interferon response. eLife. 2016;5:e16220.

53. Malle L, Martin-Fernandez M, Buta S, Richardson A, Bush D, Bogunovic D. Excessive negative regulation of type I interferon disrupts viral control in individuals with Down syndrome. Immunity. 2022;55:2074–2084.e5.

54. Ge Y, Paisie TK, Chen S, Concannon P. UBASH3A Regulates the Synthesis and Dynamics of TCR–CD3 Complexes. The Journal of Immunology. 2019;203:2827–36.

55. Kuilman T, Velds A, Kemper K, Ranzani M, Bombardelli L, Hoogstraat M, et al. CopywriteR: DNA copy number detection from off-target sequence data. Genome Biol. 2015;16:49.

56. Scheinin I, Sie D, Bengtsson H, Van De Wiel MA, Olshen AB, Van Thuijl HF, et al. DNA copy number analysis of fresh and formalin-fixed specimens by shallow whole-genome sequencing with identification and exclusion of problematic regions in the genome assembly. Genome Res. 2014;24:2022–32.

57. McKenna A, Hanna M, Banks E, Sivachenko A, Cibulskis K, Kernytsky A, et al. The Genome Analysis Toolkit: A MapReduce framework for analyzing next-generation DNA sequencing data. Genome Res. 2010;20:1297–303.

58. Robinson MD, Strbenac D, Stirzaker C, Statham AL, Song J, Speed TP, et al. Copy-number-aware differential analysis of quantitative DNA sequencing data. Genome Res. 2012;22:2489–96.

59. Qiu X, Feit AS, Feiglin A, Xie Y, Kesten N, Taing L, et al. CoBRA: Containerized Bioinformatics Workflow for Reproducible ChIP/ATAC-seq Analysis. Genomics, Proteomics & Bioinformatics. 2021;19:652–61.

60. Ashoor H, Hérault A, Kamoun A, Radvanyi F, Bajic VB, Barillot E, et al. HMCan: a method for detecting chromatin modifications in cancer samples using ChIP-seq data. Bioinformatics. 2013;29:2979–86.

61. Ashoor H, Louis-Brennetot C, Janoueix-Lerosey I, Bajic VB, Boeva V. HMCan-diff: a method to detect changes in histone modifications in cells with different genetic characteristics. Nucleic Acids Res. 2017;:gkw1319.

62. Corces MR, Trevino AE, Hamilton EG, Greenside PG, Sinnott-Armstrong NA, Vesuna S, et al. An improved ATAC-seq protocol reduces background and enables interrogation of frozen tissues. 2017;:15.

63. Picelli S, Sandberg R. Tn5 transposase and tagmentation procedures for massively scaled sequencing projects.: 9.

64. Martin M. Cutadapt removes adapter sequences from high-throughput sequencing reads. EMBnet j. 2011;17:10.

65. Zhang Y, Liu T, Meyer CA, Eeckhoute J, Johnson DS, Bernstein BE, et al. Model-based Analysis of ChIP-Seq (MACS). Genome Biol. 2008;9:R137.

66. Li Q, Brown JB, Huang H, Bickel PJ. Measuring reproducibility of high-throughput experiments. Ann Appl Stat. 2011;5.

67. Robinson MD, McCarthy DJ, Smyth GK. edgeR: a Bioconductor package for differential expression analysis of digital gene expression data. Bioinformatics. 2010;26:139–40.

