## Supplemental Data 1 - CNR per chromosme for "Copy number normalization distinguishes differential signals driven by copy number differences in ATAC-seq and ChIP-seq"

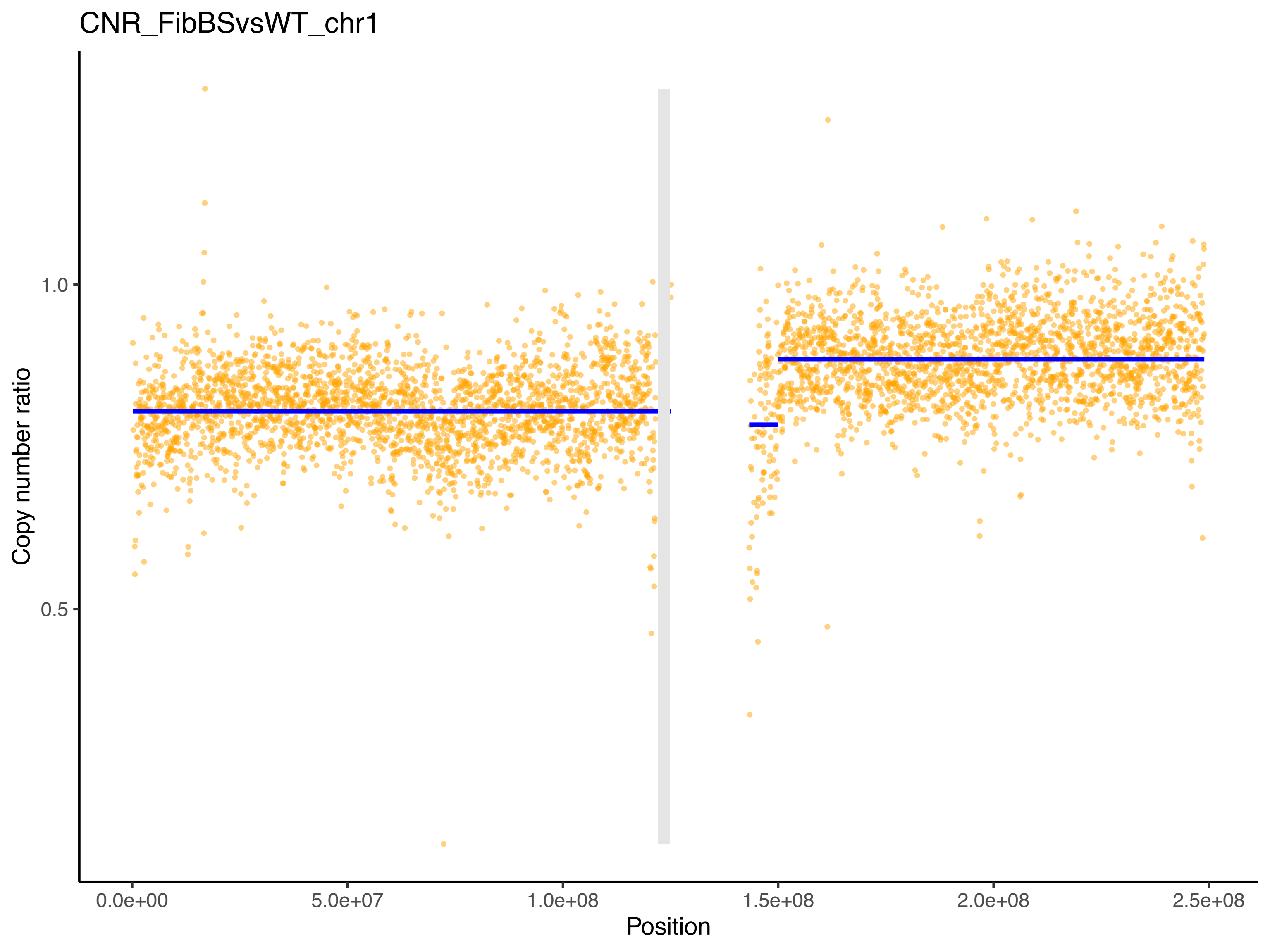
CNR_FibBSvsWT_chr1
Copy number ratio
1.0
0.5
0.0e+00
5.0e+07
1.0e+08
1.5e+08
2.0e+08
2.5e+08
Position

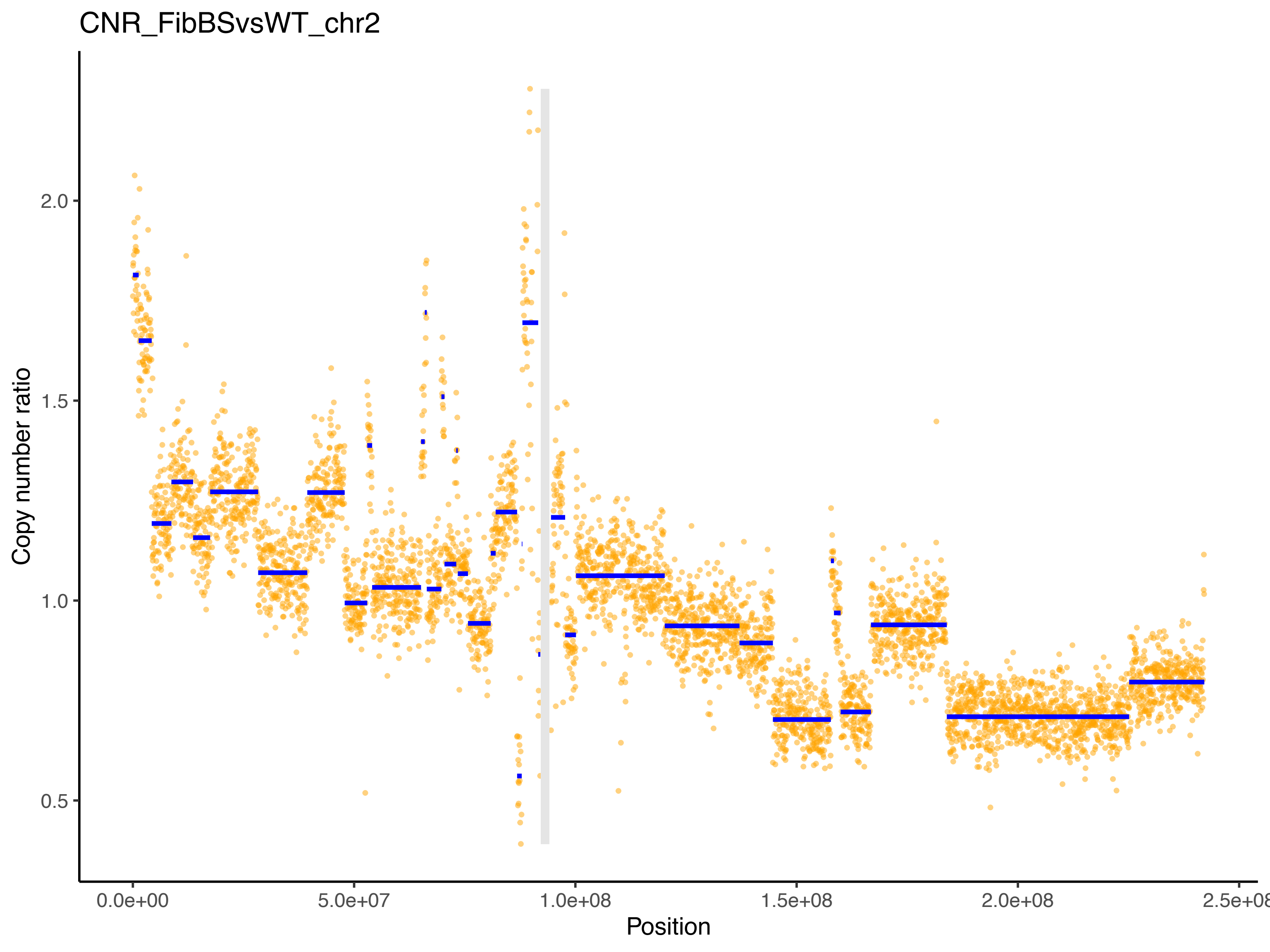

CNR_FibBSvsWT_chr2
Copy number ratio
2.0
1.5
1.0
0.5
0.0e+00
5.0e+07
1.0e+08
1.5e+08
2.0e+08
2.5e+08
Position

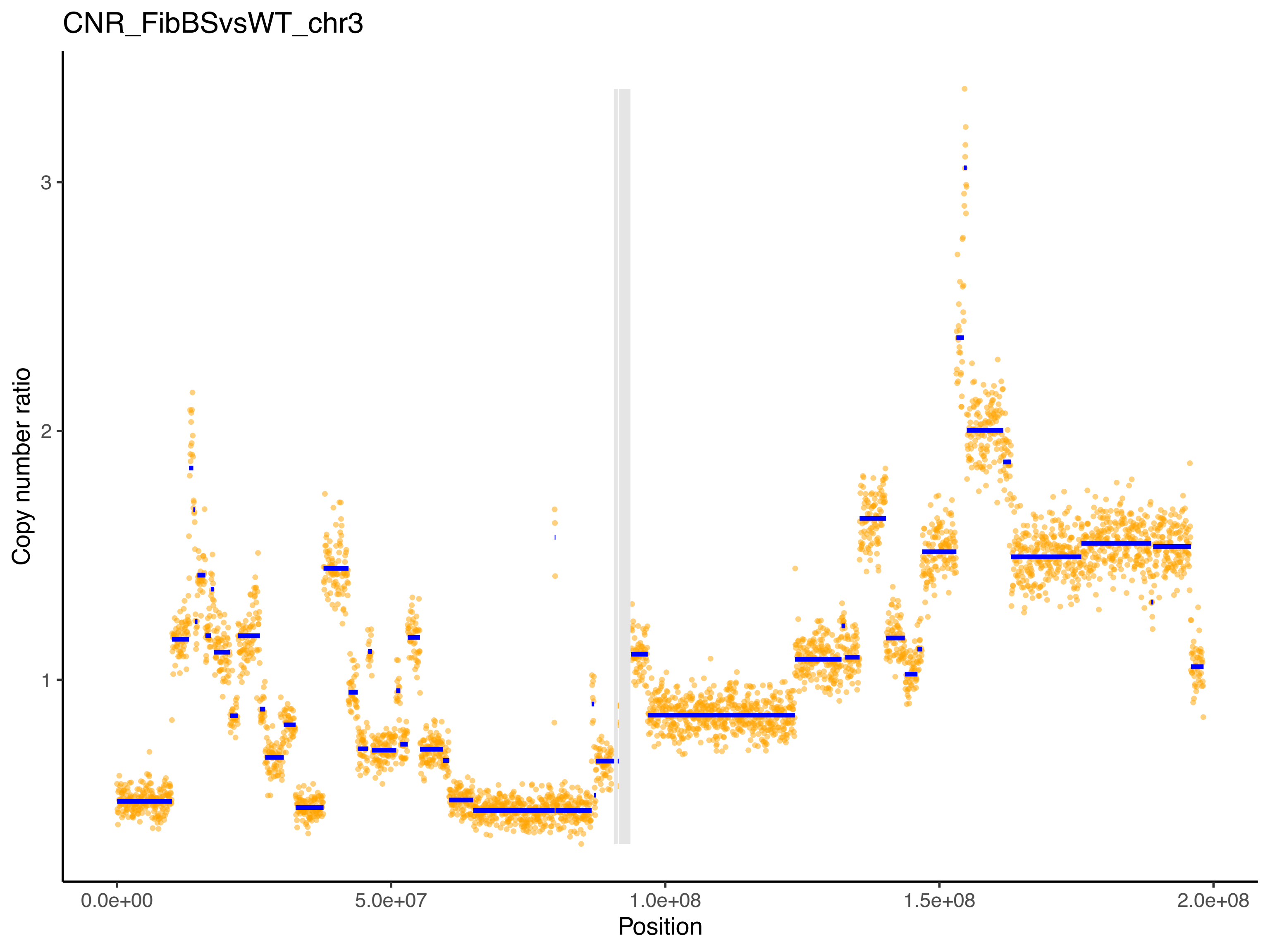

CNR_FibBSvsWT_chr3
Copy number ratio
3
2
1
0.0e+00
5.0e+07
1.0e+08
1.5e+08
2.0e+08
Position

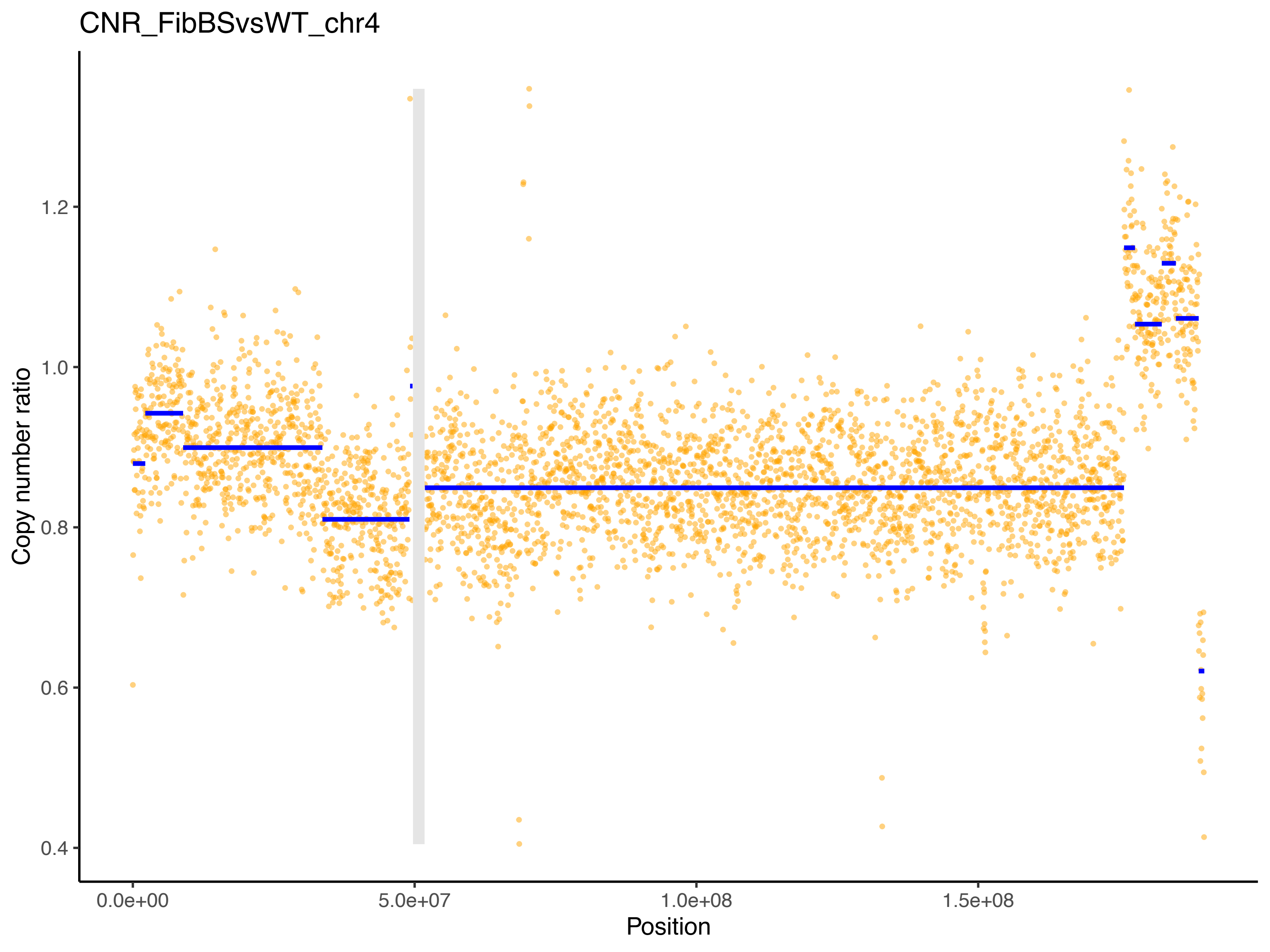

CNR_FibBSvsWT_chr4
Copy number ratio
1.2
1.0
0.8
0.6
0.4
0.0e+00
5.0e+07
1.0e+08
1.5e+08
Position

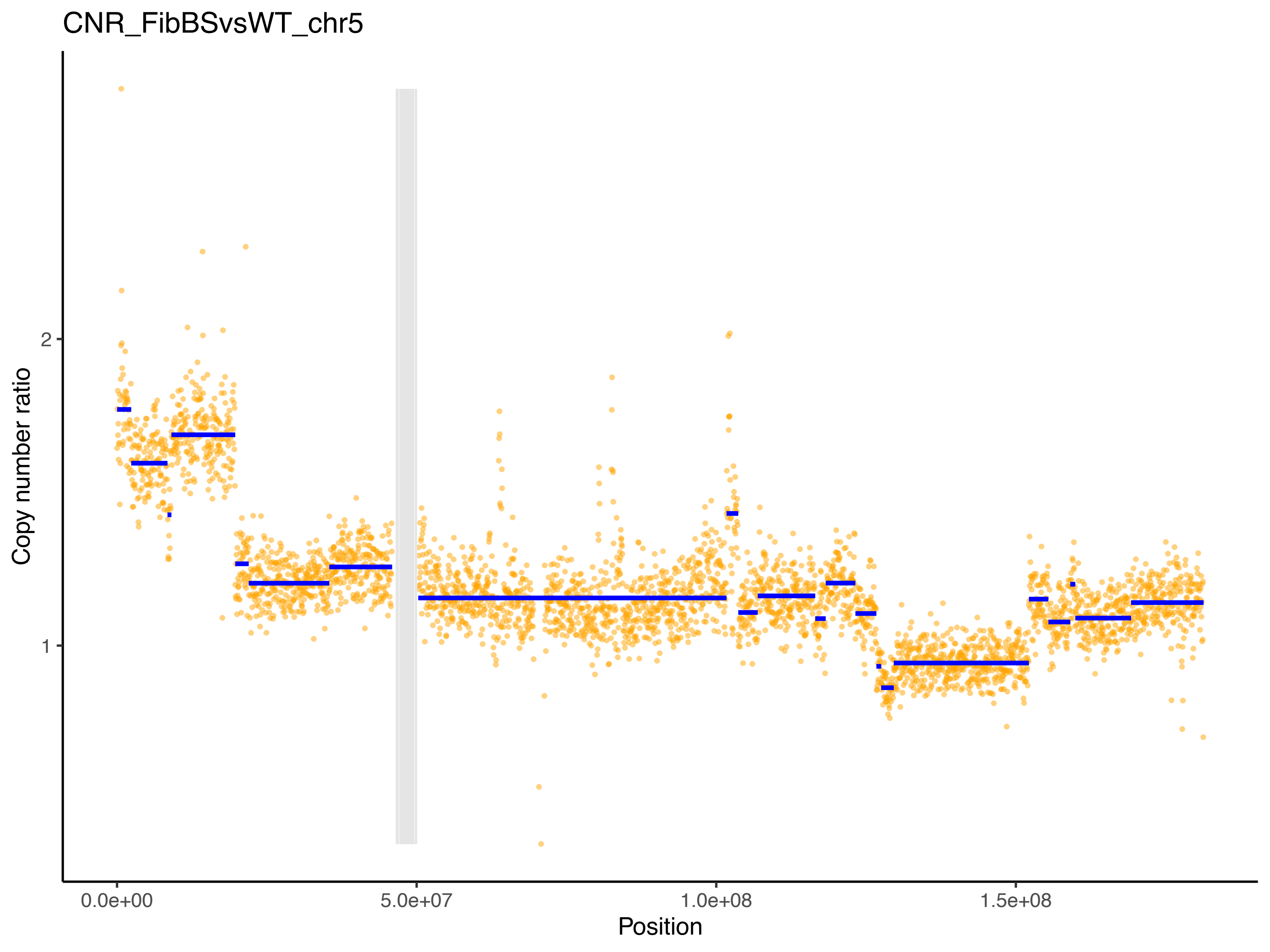

CNR_FibBSvsWT_chr5
Copy number ratio
2
1
0.0e+00
5.0e+07
1.0e+08
1.5e+08
Position

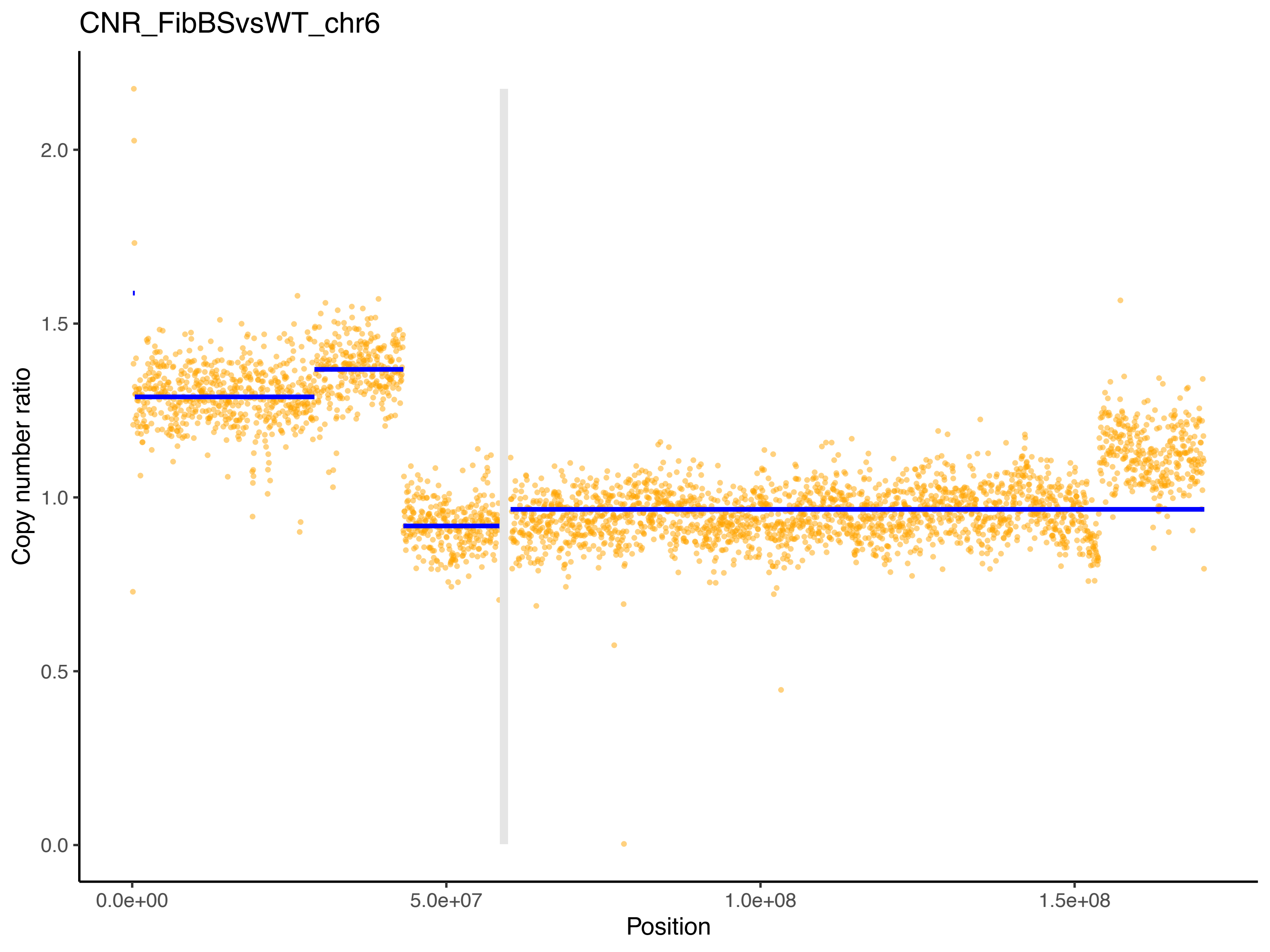
CNR_FibBSvsWT_chr6
Copy number ratio
0.0
0.5
1.0
1.5
2.0
0.0e+00
5.0e+07
1.0e+08
1.5e+08
Position

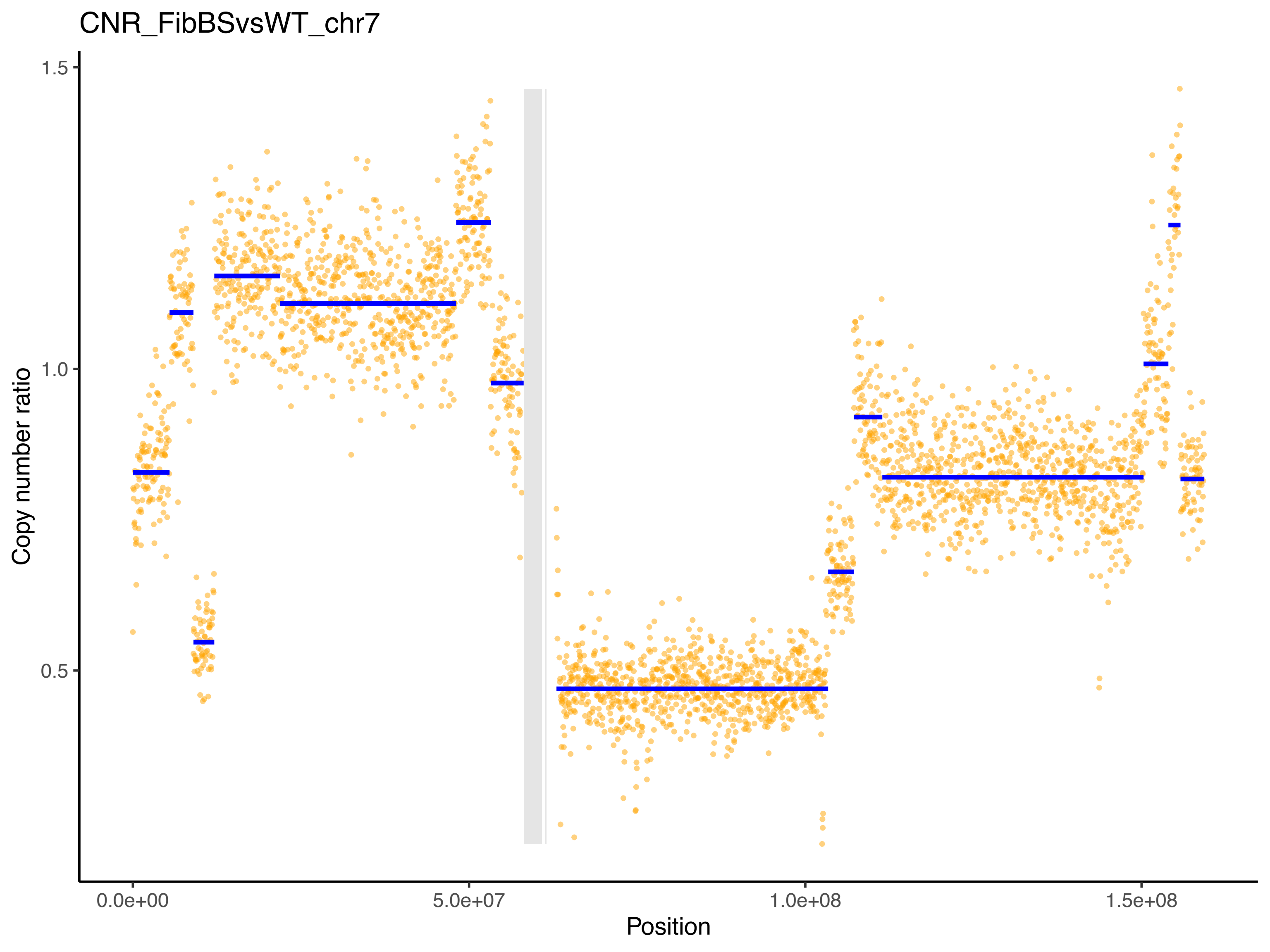

CNR_FibBSvsWT_chr7
Copy number ratio
1.5
1.0
0.5
0.0e+00
5.0e+07
1.0e+08
1.5e+08
Position

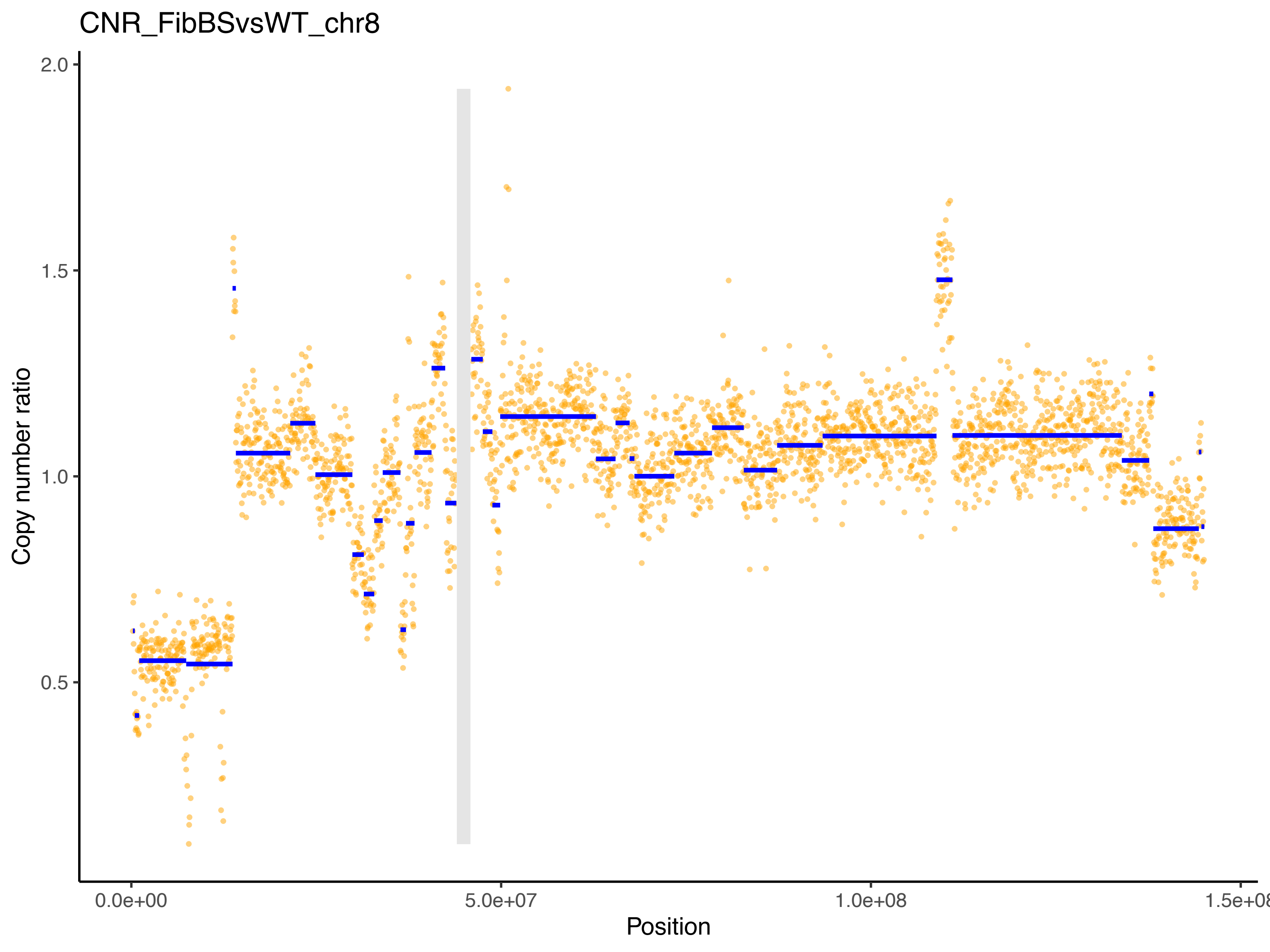
CNR_FibBSvsWT_chr8
Copy number ratio
2.0
1.5
1.0
0.5
0.0e+00
5.0e+07
1.0e+08
1.5e+08
Position

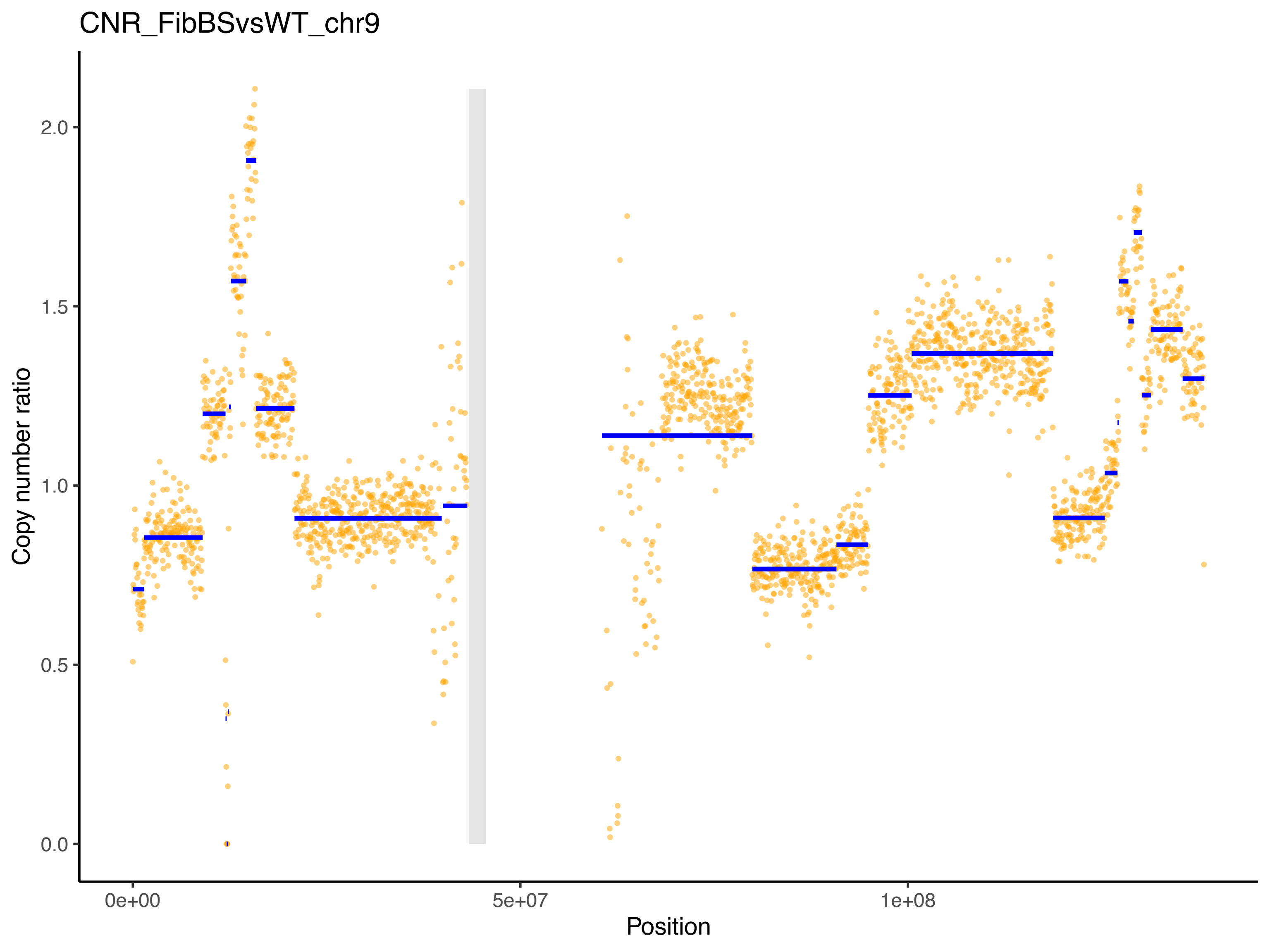
CNR_FibBSvsWT_chr9
Copy number ratio
0.0
0.5
1.0
1.5
2.0
0e+00
5e+07
1e+08
Position

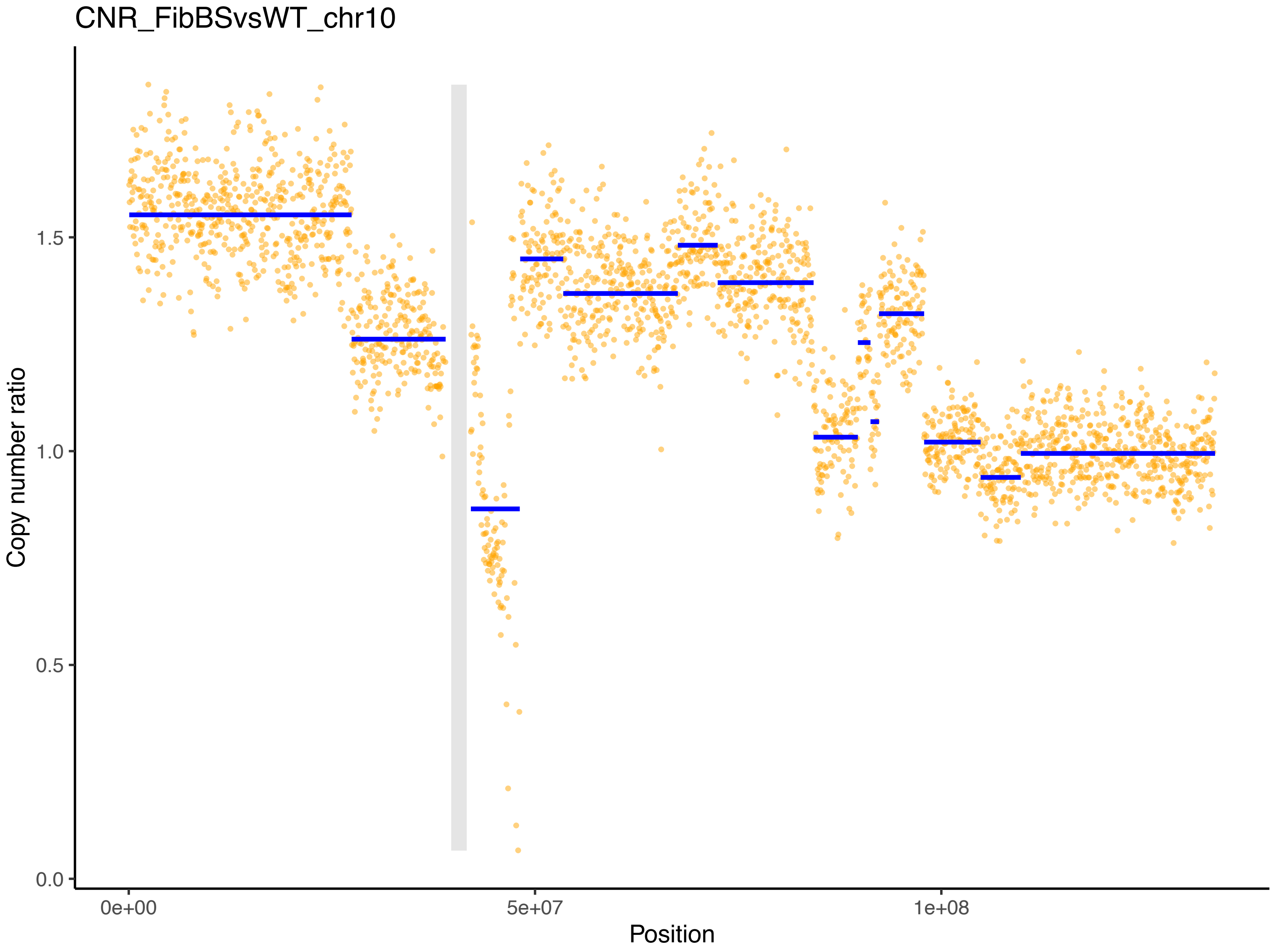

CNR_FibBSvsWT_chr10
Copy number ratio
1.5
1.0
0.5
0.0
0e+00
5e+07
1e+08
Position

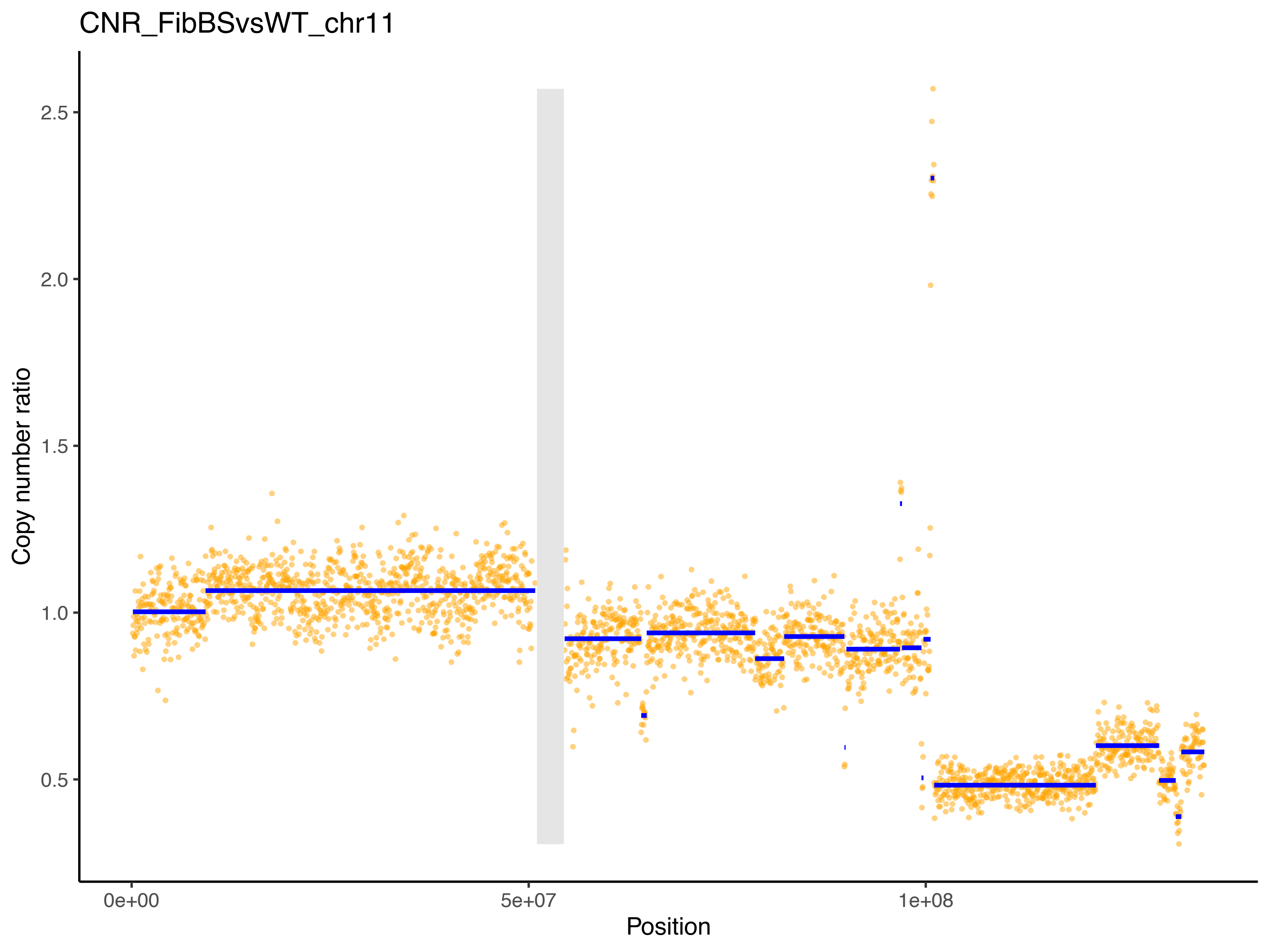

CNR_FibBSvsWT_chr11
Copy number ratio
2.5
2.0
1.5
1.0
0.5
0e+00
5e+07
1e+08
Position

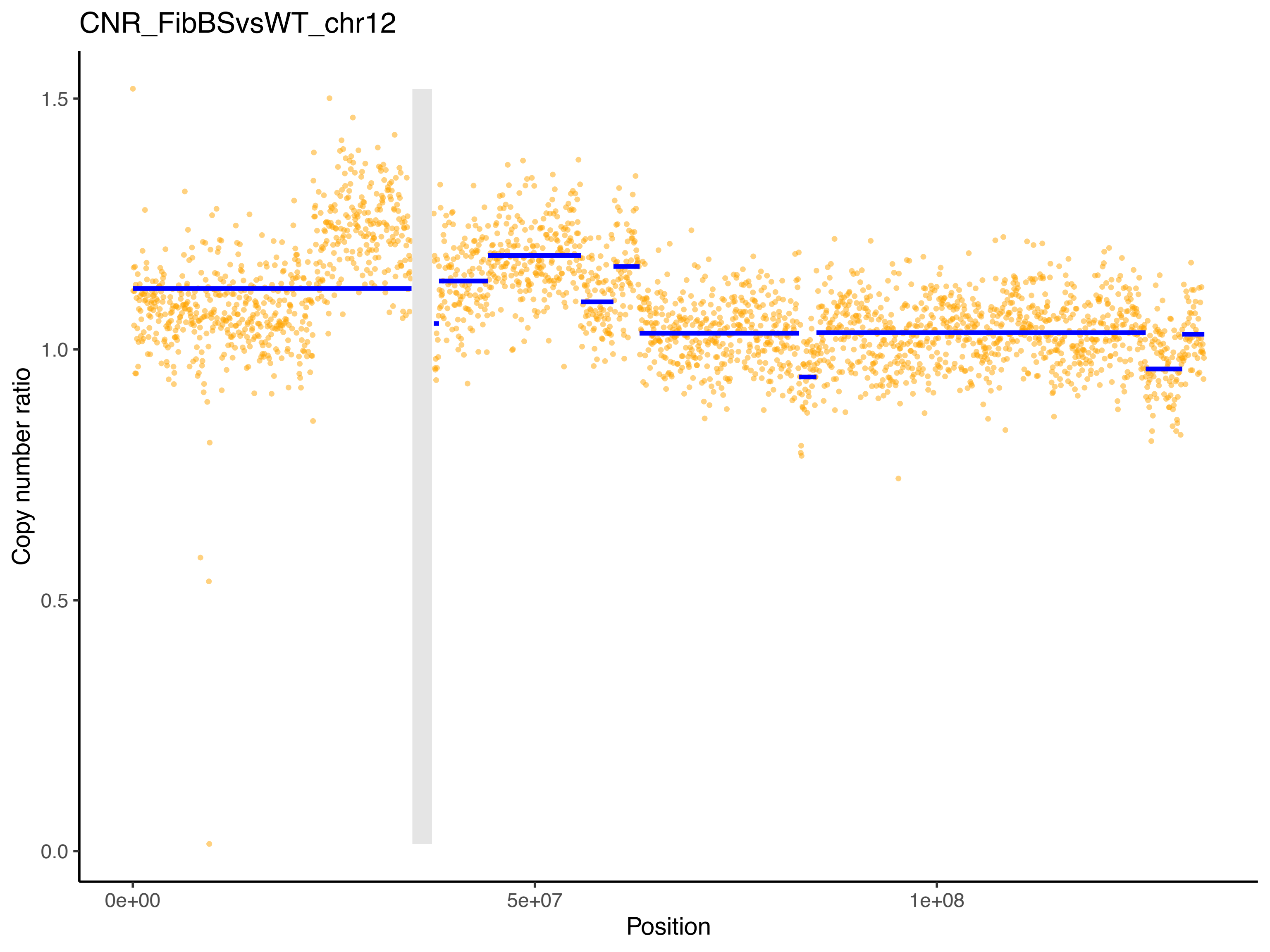
CNR_FibBSvsWT_chr12
Copy number ratio
1.5
1.0
0.5
0.0
0e+00
5e+07
1e+08
Position

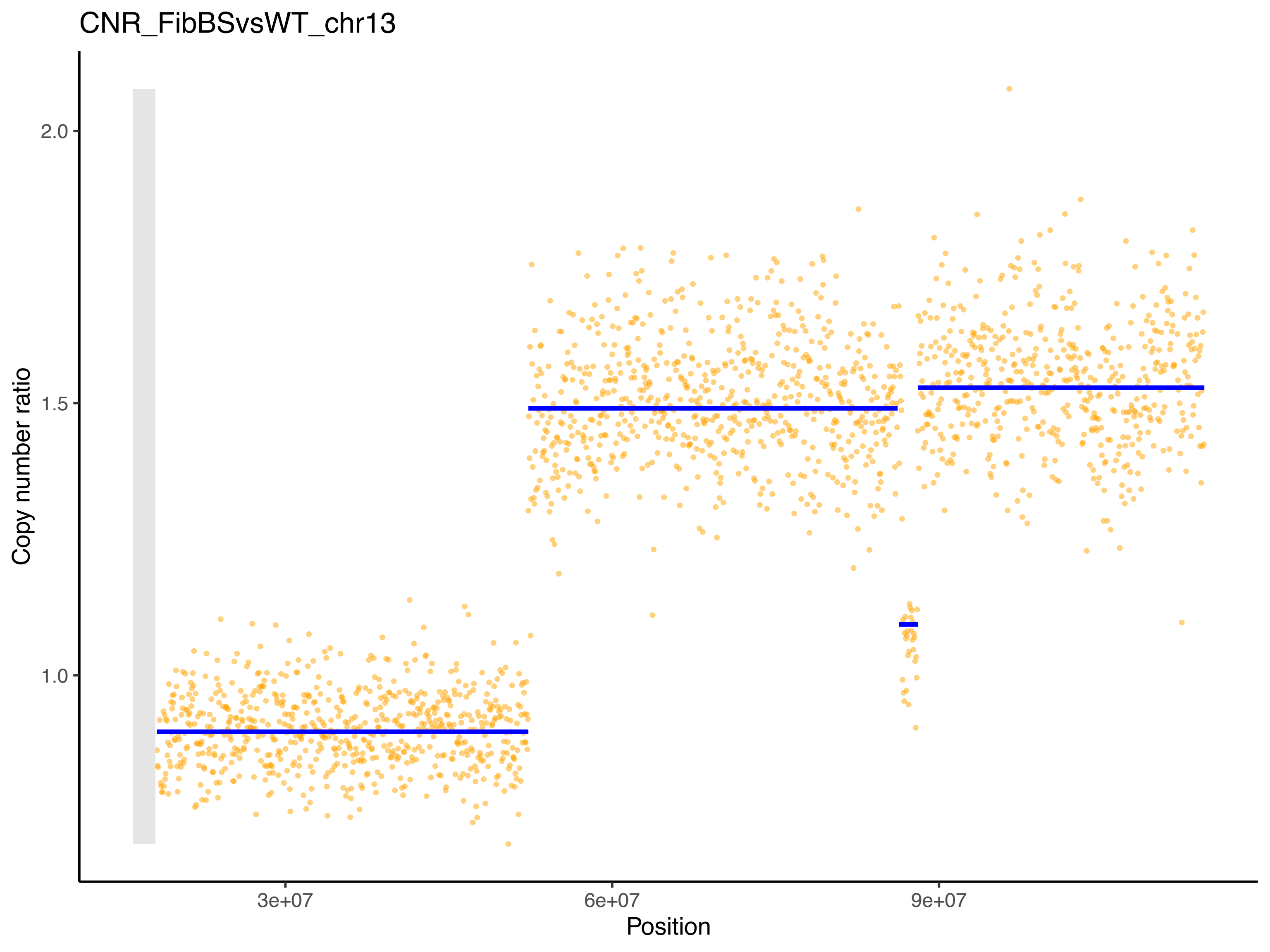

CNR_FibBSvsWT_chr13
Copy number ratio
2.0
1.5
1.0
3e+07
6e+07
9e+07
Position

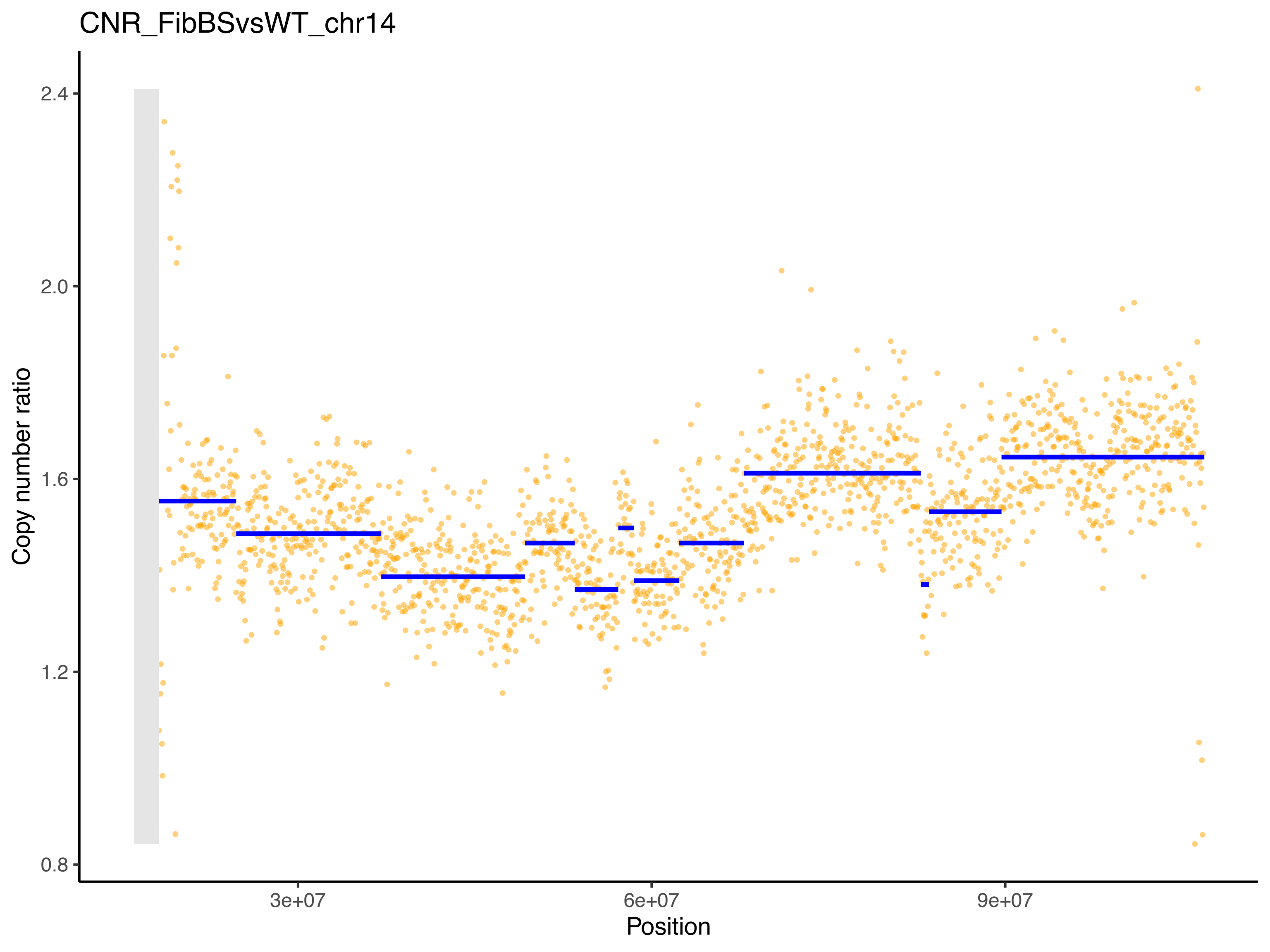

CNR_FibBSvsWT_chr14
Copy number ratio
2.4
2.0
1.6
1.2
0.8
3e+07
6e+07
9e+07
Position

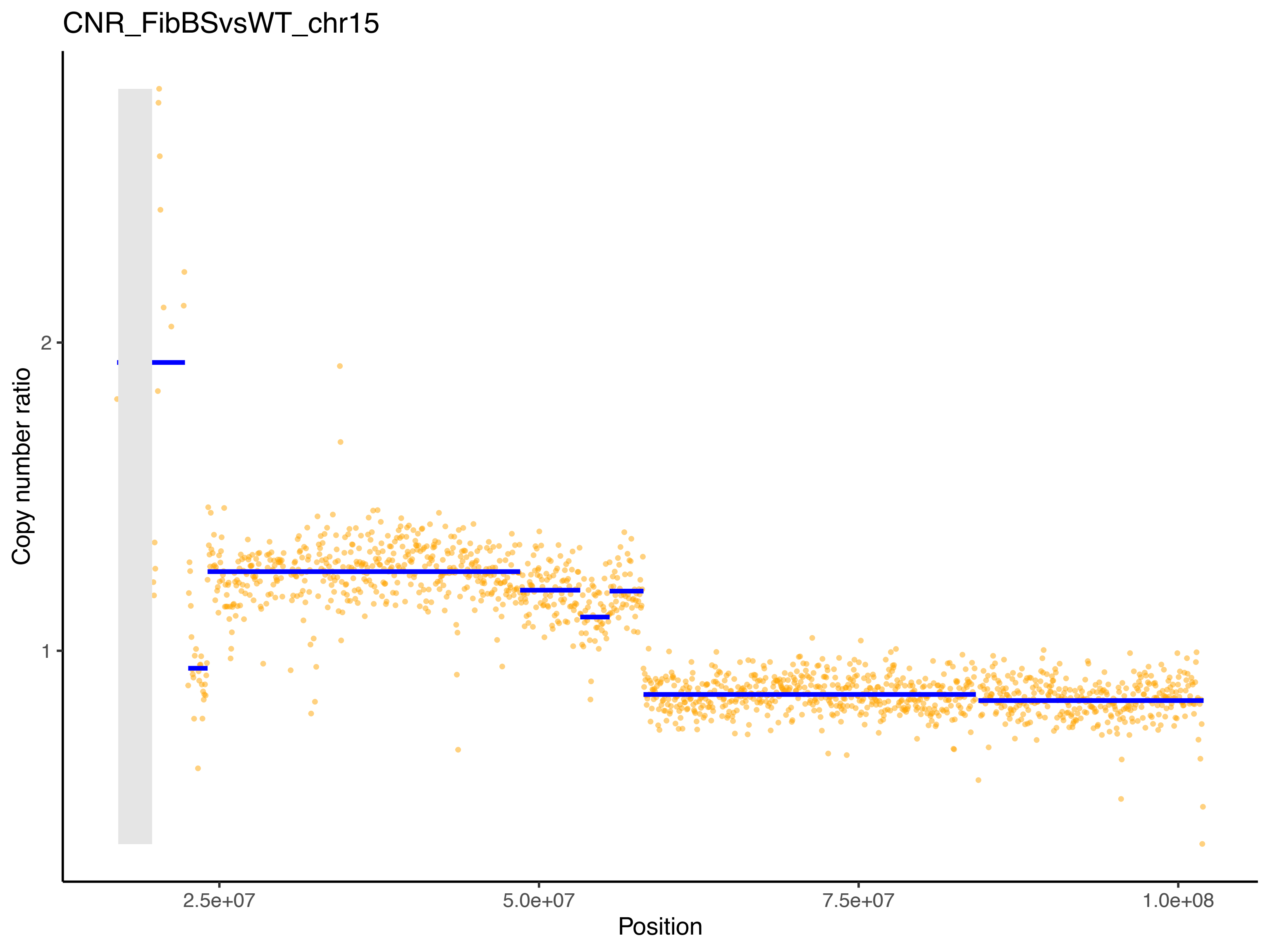
CNR_FibBSvsWT_chr15
Copy number ratio
2
1
2.5e+07
5.0e+07
7.5e+07
1.0e+08
Position

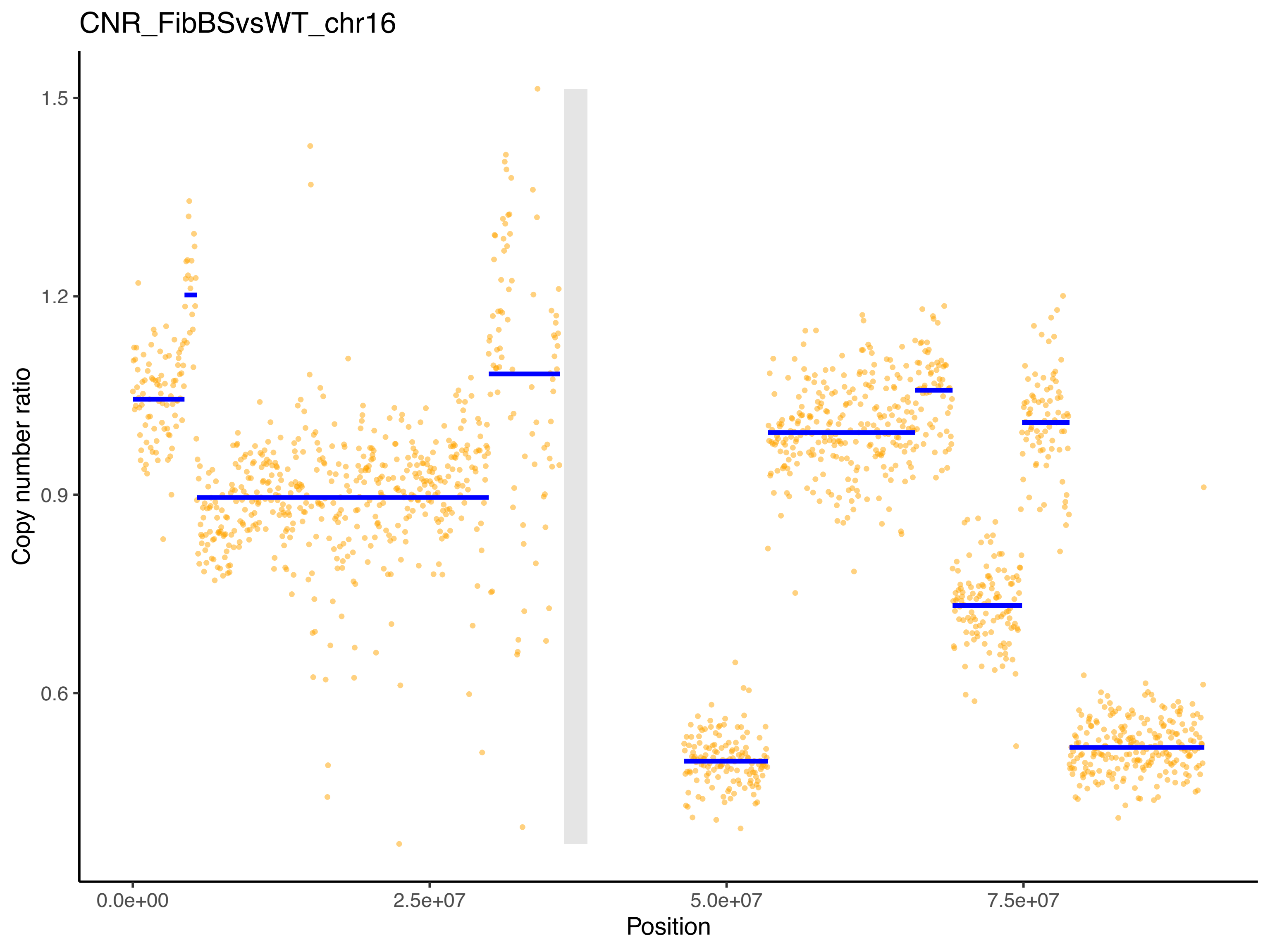

CNR_FibBSvsWT_chr16
Copy number ratio
1.5
1.2
0.9
0.6
0.0e+00
2.5e+07
5.0e+07
7.5e+07
Position

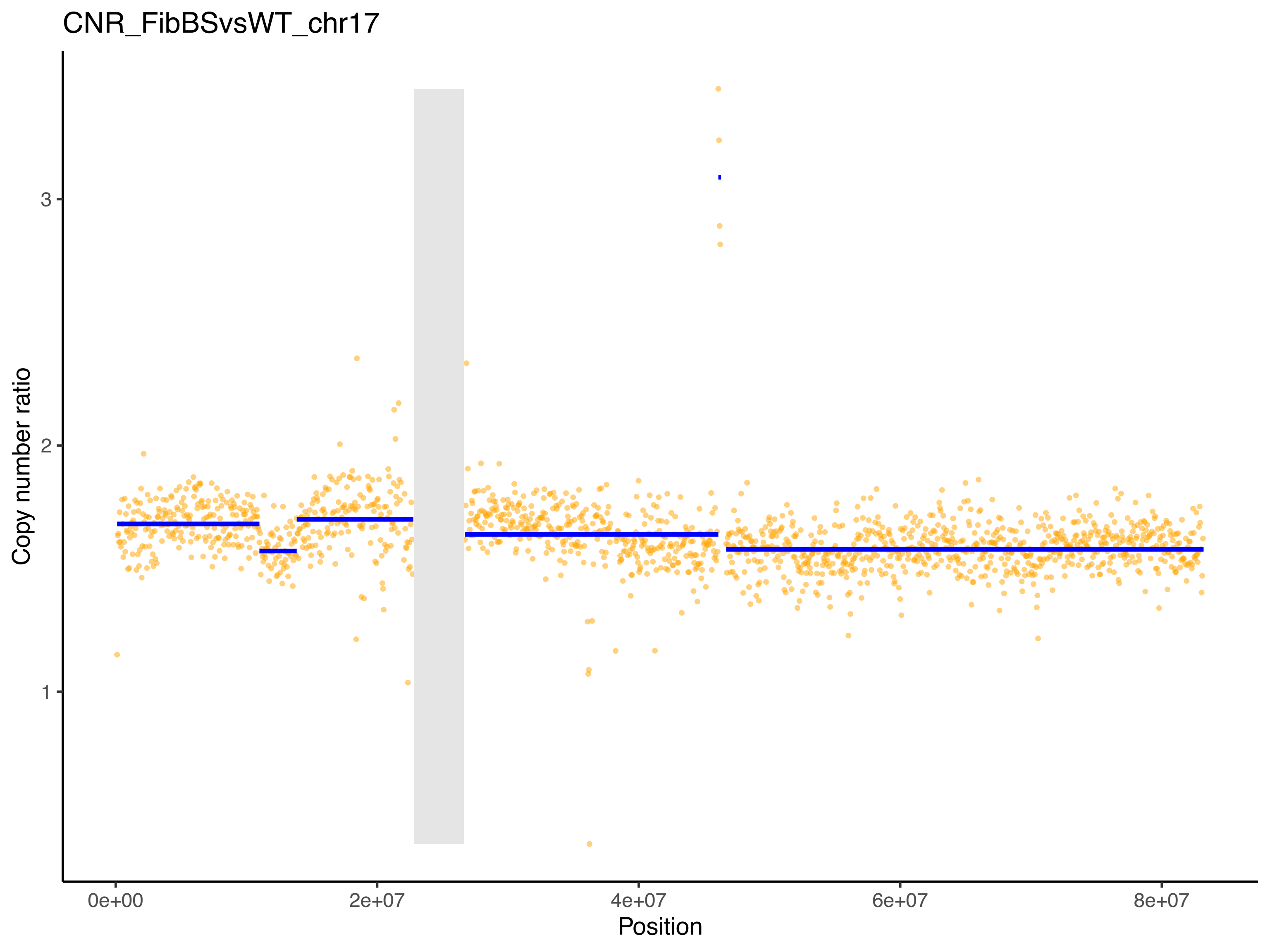
CNR_FibBSvsWT_chr17
Copy number ratio
3
2
1
0e+00
2e+07
4e+07
6e+07
8e+07
Position

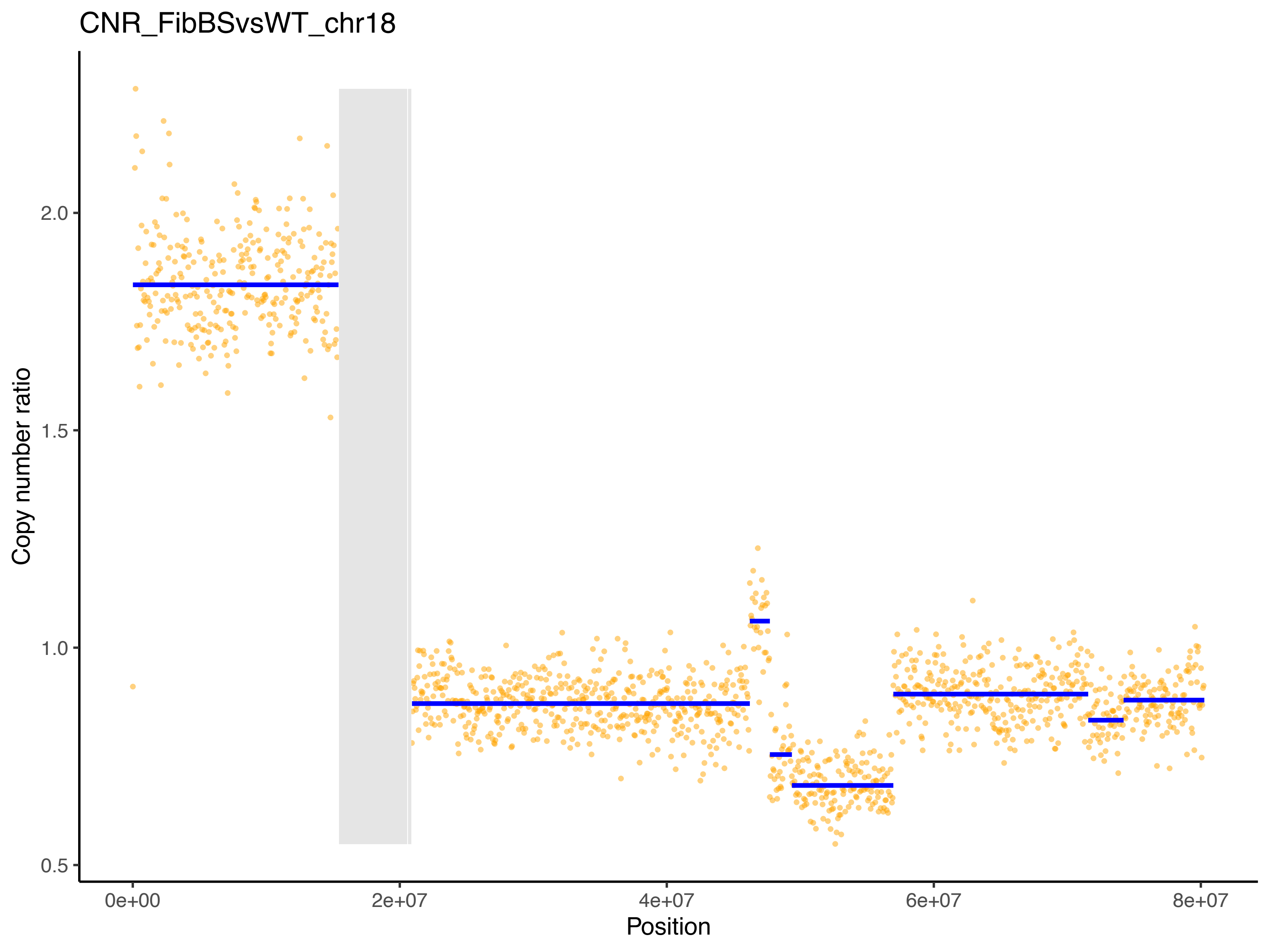

CNR_FibBSvsWT_chr18
Copy number ratio
2.0
1.5
1.0
0.5
0e+00
2e+07
4e+07
6e+07
8e+07
Position

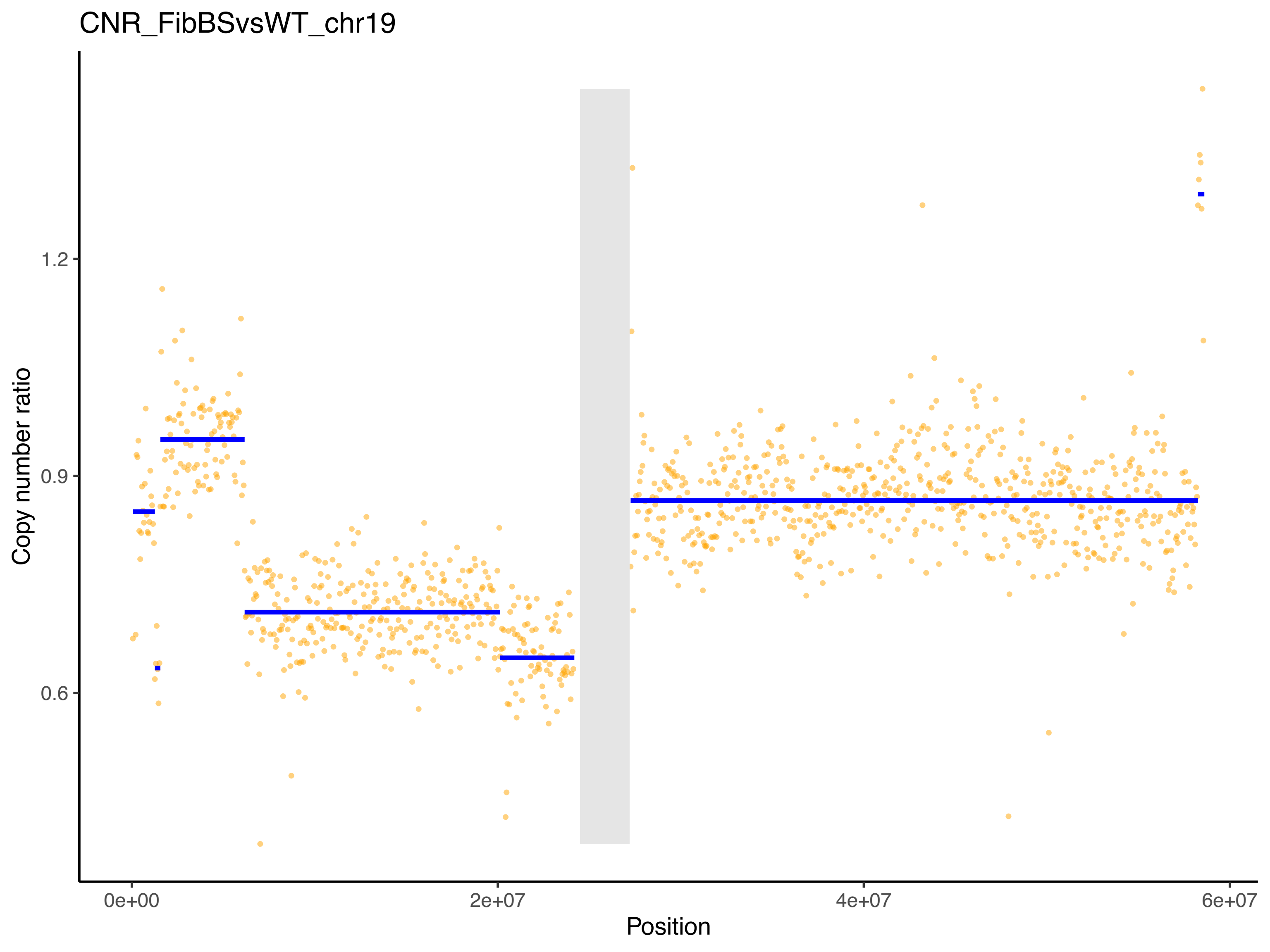
CNR_FibBSvsWT_chr19
Copy number ratio
1.2
0.9
0.6
0e+00
2e+07
4e+07
6e+07
Position

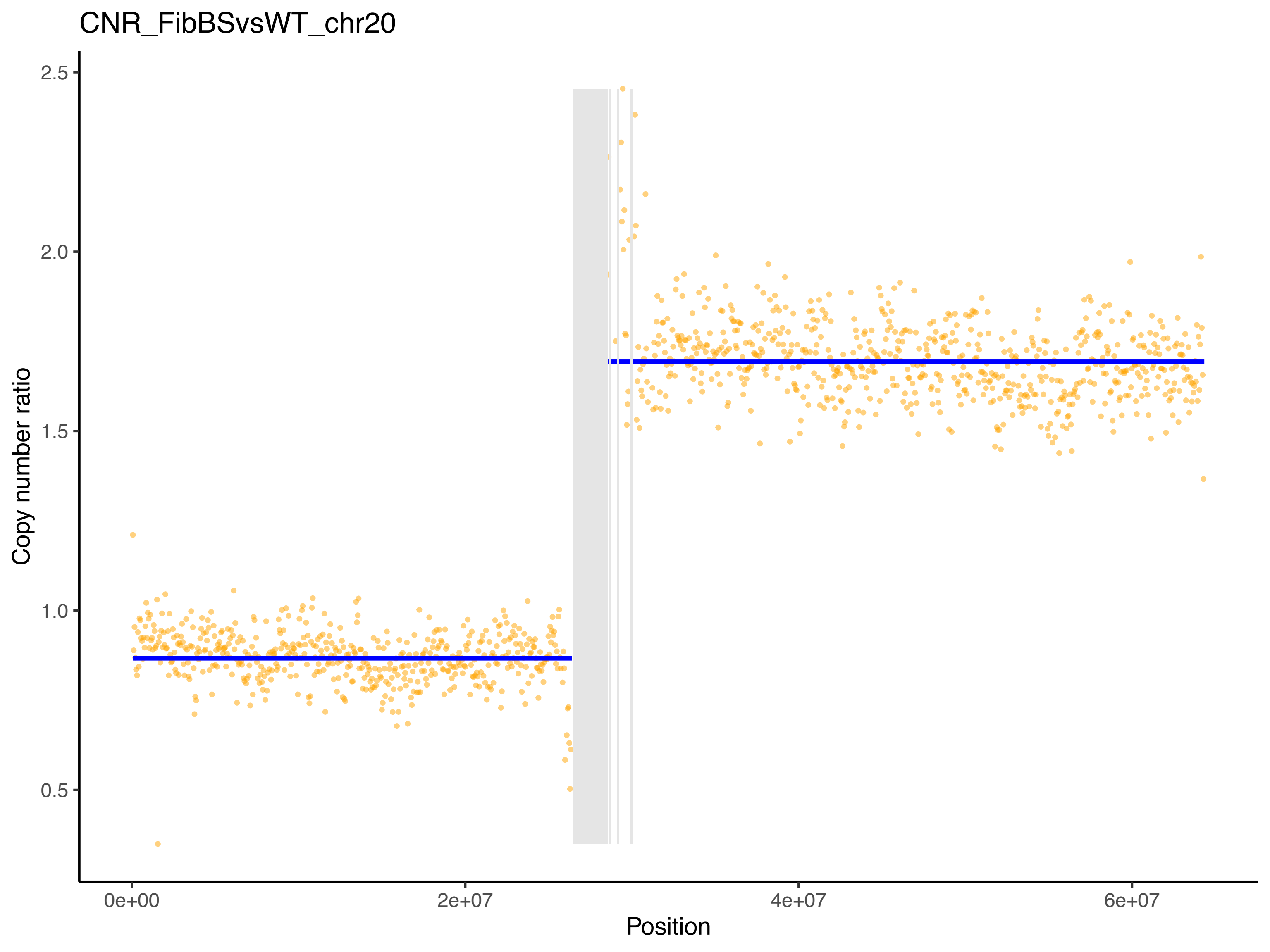
CNR_FibBSvsWT_chr20
Copy number ratio
2.5
2.0
1.5
1.0
0.5
0e+00
2e+07
4e+07
6e+07
Position

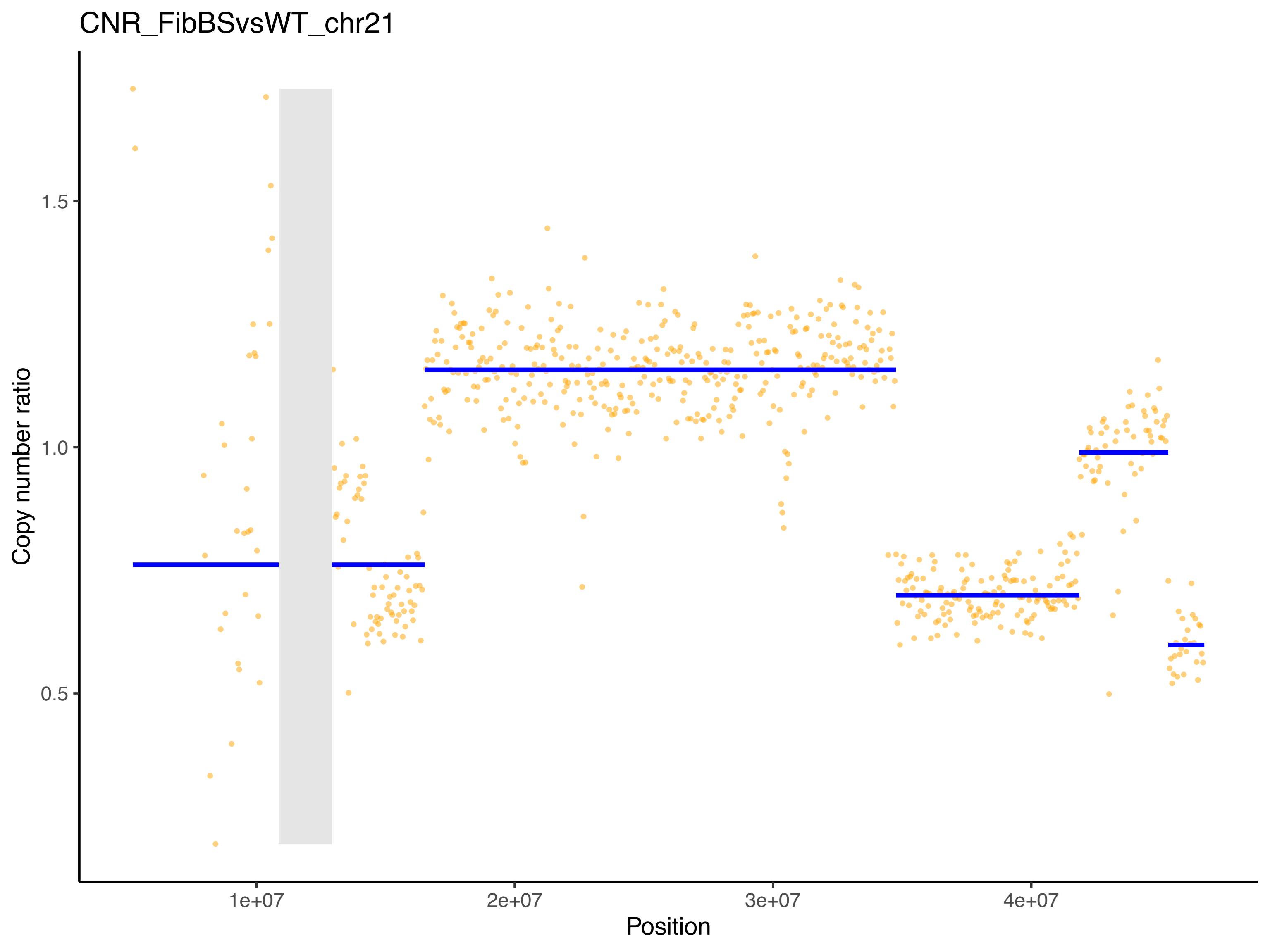
CNR_FibBSvsWT_chr21
Copy number ratio
1.5
1.0
0.5
1e+07
2e+07
3e+07
4e+07
Position

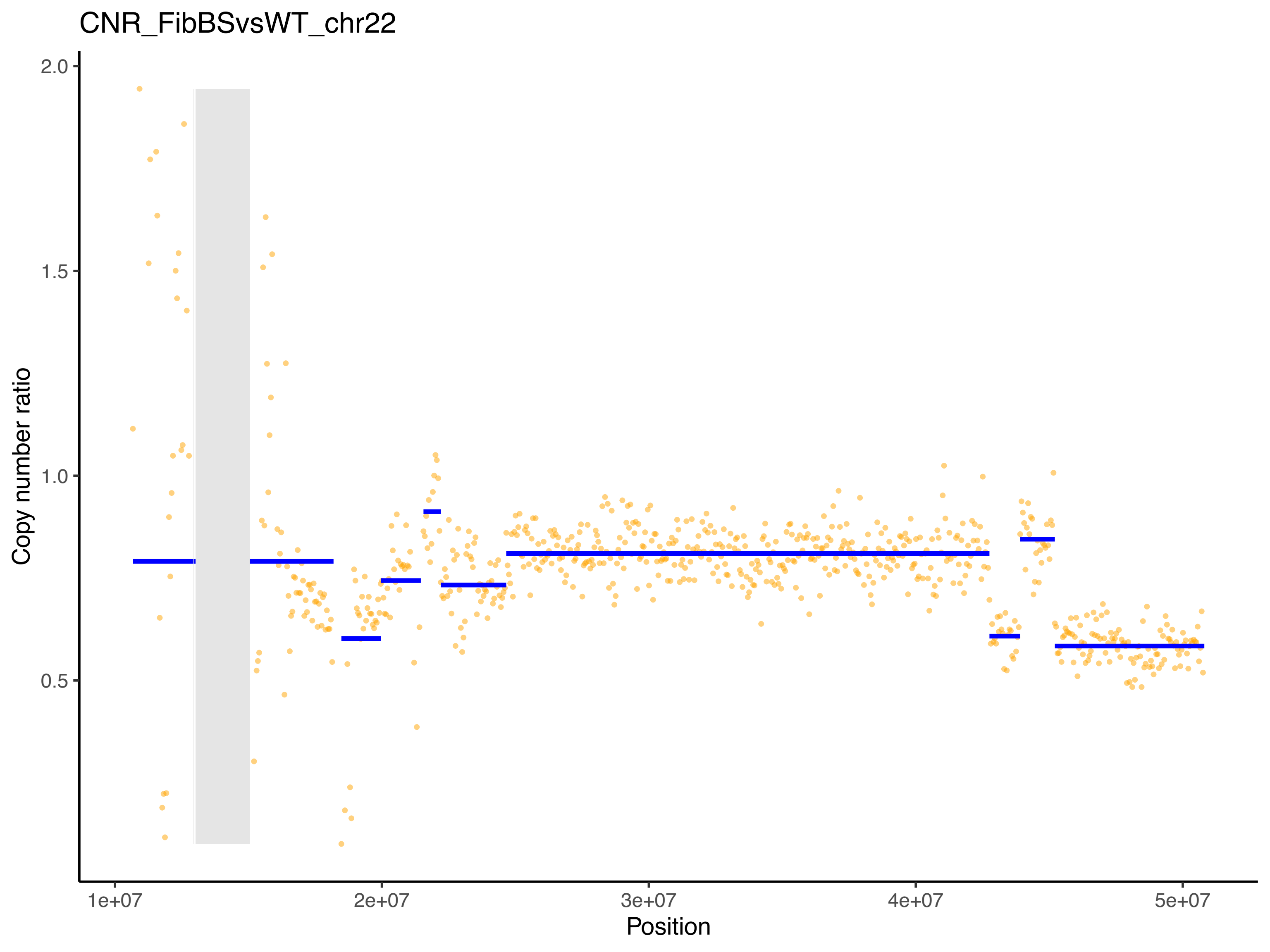
CNR_FibBSvsWT_chr22
Copy number ratio
2.0
1.5
1.0
0.5
1e+07
2e+07
3e+07
4e+07
5e+07
Position

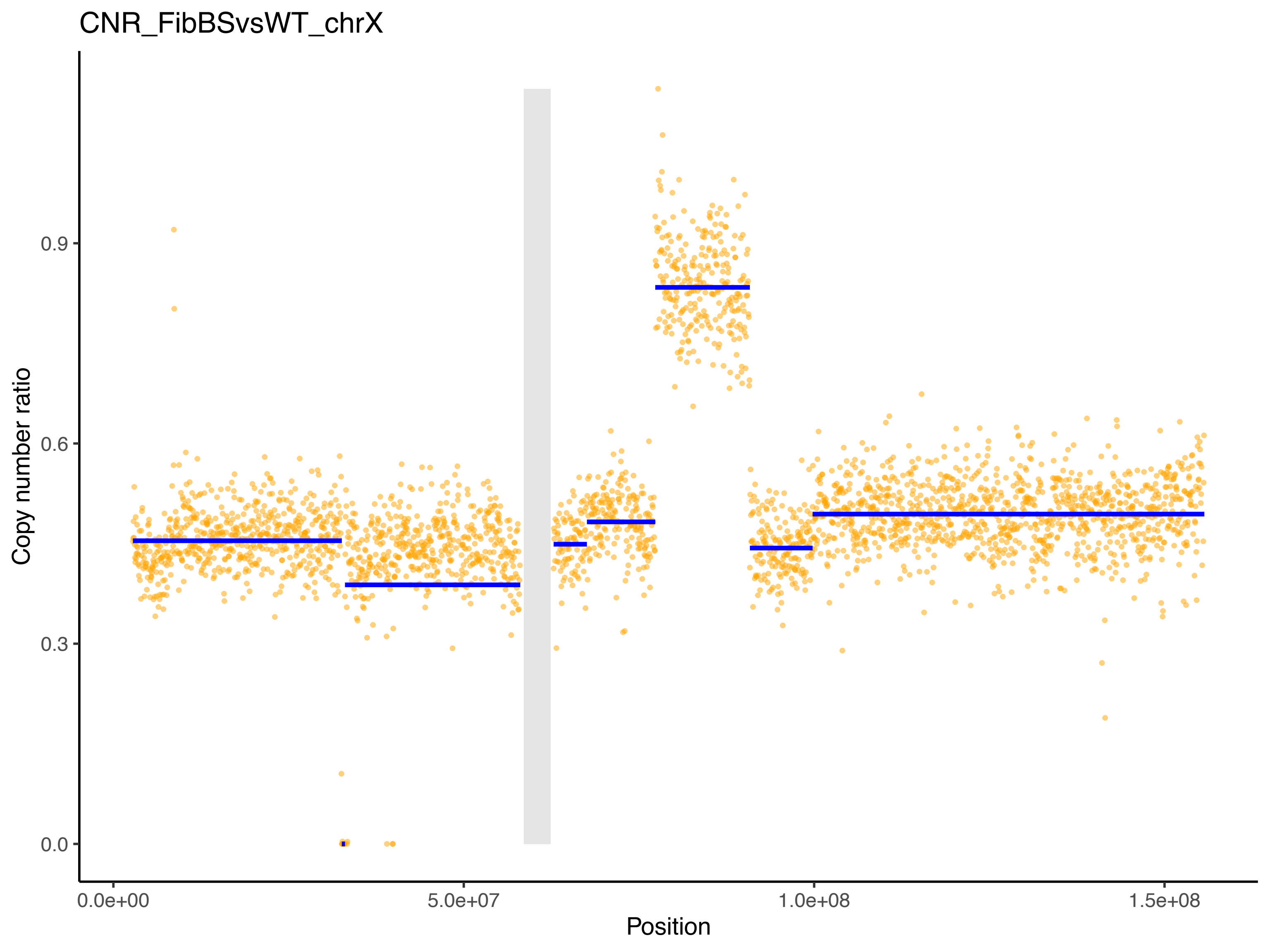
CNR_FibBSvsWT_chrX
Copy number ratio
0.0
0.3
0.6
0.9
0.0e+00
5.0e+07
1.0e+08
1.5e+08
Position
