## Supplementary Data 2 - DifferentialAnalysis CNV pipeline for "Copy number normalization distinguishes differential signals driven by copy number differences in ATAC-seq and ChIP-seq"

### Differential analysis of ATACseq with/out copy number normalization

#### pre-alignment

##### trim reads with cutadapt

```
if [ ! -e "/tmp/dsu" ];then mkdir /tmp/dsu;fi

unset PYTHONPATH
export LD_LIBRARY_PATH=
export PYTHONPATH=lib/python3.8/site-packages:$PYTHONPATH


file=$1;
echo $file

fbname=$(basename $file _R1_001.fastq.gz)
echo $fbname

dir=$(dirname $file)
echo $dir

#- O 10: minimal overlap witht he specified adapter sequence is 10nt
#-m 20: keep reads longer than 20bp after trimming

#Tn5 adaptors in the following command fro ATACseq data
cutadapt --times 2 -a CTGTCTCTTATACACATCT -g AGATGTGTATAAGAGACAG -A
        CTGTCTCTTATACACATCT -G AGATGTGTATAAGAGACAG --cores=15 -O 10  -
        -nextseq-trim=15 -m 20 --pair-filter any -o
        /tmp/dsu/"$fbname"_R1_001.cutadapt.fastq.filter.gz -p
        /tmp/dsu/"$fbname"_R2_001.cutadapt.fastq.filter.gz
        $dir/"$fbname"_R1_001.fastq.gz $dir/"$fbname"_R2_001.fastq.gz
        --too-short-output=/tmp/dsu/"$fbname"_R1_001.fastq.tooshort.gz
        --too-short-paired-
        output=/tmp/dsu/"$fbname"_R2_001.fastq.tooshort.gz 1>
        "$fbname".cutadapt.err 2> "$fbname".cutadapt.out

# Truseq adaptors in the following command for ChIPseq data
cutadapt --times 2 -a AGATCGGAAGAGCACACGTCTGAACTCCAGTCAC -g
        ACACTCTTTCCCTACACGACGCTCTTCCGATCT -A
        AGATCGGAAGAGCGTCGTGTAGGGAAAGAGTGTG -G
        GTGACTGGAGTTCAGACGTGTGCTCTTCCGATCT --cores=8 -O 10 --nextseq-
        trim=15 -m 22 --pair-filter=any -o
        /tmp/dsu/"$fbname"_R1_001.cutadapt.fastq.filter.gz -p
        /tmp/dsu/"$fbname"_R2_001.cutadapt.fastq.filter.gz
        $dir/"$fbname"_R1_001.fastq.gz $dir/"$fbname"_R2_001.fastq.gz
        --too-short-output=$dir/"$fbname"_R1_001.fastq.small.gz --too-
        short-paired-output=$dir/"$fbname"_R2_001.fastq.small.gz

echo "Cutadapt is finished"
```

#### reads alignment and filtering

##### align reads

```
bwa mem -t 10 /fml/chones/genome/gbdb/hg38/hg38.fa \
    /tmp/dsu/"$fbname"_R1_001.cutadapt.fastq.filter.gz
        /tmp/dsu/"$fbname"_R2_001.cutadapt.fastq.filter.gz \
    -R "@RG\tID:$fbname\tSM:$fbname\tLB:ChIPseq\tPL:NovaseqS4.2x150"
        |\
    samtools view -bh -@ 10 - > /tmp/dsu/$fbname.hg38.bam

samtools sort -@ 10 -l 9 \
    -T /tmp/dsu/$fbname.tmpsort \
    -o /tmp/dsu/$fbname.hg38.sorted.bam /tmp/dsu/$fbname.hg38.bam
mv /tmp/dsu/"$fbname"_R?_001.cutadapt.fastq.filter.gz $dir/
mv /tmp/dsu/$fbname.hg38.sorted.bam $dir/
rm /tmp/dsu/$fbname.hg38.bam

echo "Mapping to hg38 of sample "$file" is finished"
```

##### filter reads

```
#mark duplicates
/fml/chones/data/Dingwen/SCRIPTS/Picard_MarkDuplicate.cluster.sh $file

#!/bin/bash
file=$1
fbname=$(basename $file .sorted.bam)
echo $fbname
```

```
dir=$(dirname $file)
echo $dir

if [ ! -e "/tmp/dsu" ]
                         then mkdir /tmp/dsu
fi

java -Xmx3g -XX:ParallelGCThreads=2 -Djava.io.tmpdir=/tmp/dsu -jar
        picard-2.18.25/picard.jar MarkDuplicates \
    I=$file \
    O=/tmp/dsu/$fbname.pMarkdup.bam \
    M=/tmp/dsu/$fbname.pMarkdup.metrics\
    REMOVE_DUPLICATES=false CREATE_INDEX=true
        VALIDATION_STRINGENCY=STRICT

# remove reads with low quality
#Filter out Mitochondrial reads, reads with mapping quality lower than
        20 with samtools

file=$1
echo $file
fbname=$(basename $file .pMarkdup.bam)
echo $fbname
dir=$(dirname $file)
echo $dir

if [ ! -e "/tmp/dsu" ]
                         then mkdir /tmp/dsu
fi

samtools view -@ 3 -h -F 3588 -q 20 $file |grep -v "chrM"| \
    samtools view -bh -@ 3 - >/tmp/dsu/$fbname.deDupMtq20.bam
samtools index -@ 3 /tmp/dsu/$fbname.deDupMtq20.bam

#filter out reads in hg38 blacklist regions
#there are 2 black list, one form UCSC:
        /fml/mickle/data/Dingwen/HiSeq3000/hg38.blacklist.bed
#hg38 blacklist downloaded from Ensemble:
        #/fml/mickle/data/Dingwen/HiSeq3000/Coriell_G4ChIP/ENCODE_Grh38_ExclusionList_ENCFF356LFX.Pos
bedtools intersect -v -abam /tmp/dsu/$fbname.deDupMtq20.bam \
     -b ENCODE_Grh38_ExclusionList_ENCFF356LFX.PositionMinus1.bed -f
        0.5 >\
     /tmp/dsu/$fbname.q20DeDupExcludableRegion.bam
```

##### peak calling

```
#subsample 20Mil reads from each sample
file=$1;
echo $file

fbname=$(basename $file .q20DeDupExcludableRegion.bam)
echo $fbname

dir=$(dirname $file)
echo $dir

#calculate the proportion for 20Mil reads (not fragments)
frac=$( samtools idxstats $file | cut -f3 | awk 'BEGIN {total=0}
        {total += $1} END {frac=20000000/total;if (frac > 1) {print 1}
        else {print frac}}')

total=$(samtools idxstats $file|datamash sum 3)
echo $total
echo $frac
#frac =$(awk 'BEGIN {print $frac/$total}')
samtools view -@ 5 -bh --subsample-seed 25 --subsample $frac $file >
        /tmp/dsu/"$fbname".subsample20Mil.bam

#make the pooled bamfile for peak calling
sanmtools merge -c -p -f -@ 10 all_the_susampled.files >
        Sample.merge.bam

# call peaks with macs2
macs2 callpeak --tempdir /tmp/dsu/ -g hs \
        -t Sample.merge.bam \
        --outdir /tmp/dsu/ \
        -n "$fbname" \
        -f BAMPE --nomodel --min-length 100 -B --SPMR --cutoff-
        analysis --keep-dup all
```

##### count reads in peaks

```
#convert peaks in narrowpeak format to gff format
cat fbname_peaksnarrowPeak |grep -v chrX |grep -v chrY |awk '{split($4,a,"_");parent=$4;gsub(/[a-z]$/
        $1"\tMACS2narrowPeak\tSO:0000684\t"$2"\t"$3"\t"$7"\t"$6"\t.\tID="$4";Name=nP_"a[length(a)]"|s
        log10pValue="$8"|-log10qvalue="$9"|peak="$1":"$2"-"$3"|summit="$1":"$2+$10";Parent="parent}'
        fbname_peaksnarrowPeak.gff
```

```
#count reads in peaks with htseq count
#!/bin/bash

#this script is to use Htseq count to count reads from a bamfile in
        each ATACseq peak

#note that use global peaks called from merged bamfiles and this
        peakset has to be transformed into gff format
#where each ATACseq peak is given a feature SO:0000684, thus -t
        SO:0000684 is used in htseq command
#Check the Htseq counting mode before running the command
#check the basename for input bamfiles

#usuage: ./Script Bamfile Peak.gff Output_directory
bamfile=$1

fbname=$(basename $bamfile .q20DeDupExcludableRegion.bam)
echo $fbname
dir=$(dirname $bamfile)
echo $dir

peakfile=$2
fbname2=$(basename $peakfile _peaks.narrowPeak.gff)
echo $fbname2

OutputDir=$3

if [ ! -e "/tmp/dsu" ]
                         then mkdir /tmp/dsu

samtools view -@ 3 -h $bamfile | htseq-count -m union -t SO:0000684 -a
        10 -s no -f sam -r pos -i ID --additional-attr Name --
        nonunique none - $peakfile >
        /tmp/dsu/"$fbname"__"$fbname2".HTseq.readcount
        2>/tmp/dsu/"$fbname"__"$fbname2".HTseq.readcount.out
mv  /tmp/dsu/"$fbname"__"$fbname2".HTseq.readcount* $OutputDir

#extract the count from all samples and make a count matrix
paste BS_D1_ATAC_S7.pool.HTseq.readcount
        BS_D2_ATAC_S11.pool.HTseq.readcount
        BS_D3_ATAC_S15.pool.HTseq.readcount
        WT_K1_ATAC_S8.pool.HTseq.readcount
        WT_K2_ATAC_S12.pool.HTseq.readcount
        WT_K3_ATAC_S16.pool.HTseq.readcount |head -n-5|awk -v OFS='\t'
        '{print $1,$3,$6,$9,$12,$15,$18}' >
        ATAC_BS_WT.pool.HTseq.readcount
```

#### optional: CNV correction with CNVkit

- note that input data here are the genomic seuqencing data from 2 samples. The bamfiles were processed and filtered exactly as described before
- CNV kit: https://cnvkit.readthedocs.io/en/stable/
- CNVkit offers a handy all-in-one command to call copy number variation, which is very convenient if the reference sample has a known karyotype. By default the reference sample is diploid and inferred copy number in the sample will only be integer

##### calling Copy number ratio

###### *run the all-in-one command*

```
#doing the analysis with the whole genome
cnvkit.py batch BS_D_gDNA_S3.hg38.q20DeDupExcludableRegion.bam \
-n WT_K_gDNA_S4.hg38.q20DeDupExcludableRegion.bam \
--fasta /genome/gbdb/hg38/hg38.fa \
#specifying whole-genome anlaysis and bin size of 50kb
--method wgs --target-avg-size 50000 --targets ../hg38.access_50kb.bed
        \
--processes 10
```

- Given the complex CNV in my samples, the log2 ratio of copy number in the sample relative to reference before calling copy number was used and this ratio was converted to the original value, as the copy number ratio (CNR). the relevant file in the ouput is `BS_D_gDNA_S3.hg38.q20DeDupExcludableRegion.cns`

```
#extract the intervals of the DNA segments
#retrive the log2 ratio and calculate the original copy number ratio
        of Fib_BS/Fib_WTm
cat BS_D_gDNA_S3.hg38.q20DeDupExcludableRegion.cns|awk -v OFS='\t'
        '{print $1,$2,$3,$5,2^$5,$6,$7,$8}' >
        BS_D_ref2K.Merged_segmentCN.with_header.bed

tail -n+2 BS_D_ref2K.Merged_segmentCN.with_header.bed|awk '{print
        $1"\t"$2"\t"$3"\t"$5}'
        >BS_D_ref2K.50kb_bin.CopyNumberRatio.bed
```

**assign scaling factor to peaks**

- peaks that overlap a segment, the copy number ratio of the segment will be the scaling factor
- for peaks that do not overlap any segments, using the copy number ratio of the nearest segment as the scaling factor

```
#peaks that overlap a segment, the copy number ratio of the segment
        will be the scaling factor
bedtools closest -a Merge.ATAC_D_K.20Meach_peaks.narrowPeak.bed -b
        BS_D_ref2K.50kb_bin.CopyNumberRatio.bed -wa -wb|awk -v
        OFS='\t' '{print $1,$2,$3,$4,$5"_"$6"_"$7,$8}' >
        Merge.ATAC_D_K.peak_scaling_factor
### apply CNV correciton to readcount matrix in R as described before
```

#### differential analysis with DESeq2

- prepare a metadata information for the samples

```
# an example metadata information table
Sample_ID   condition   library
BS_D1   BSFibroblasts   ATAC
BS_D2   BSFibroblasts   ATAC
BS_D3   BSFibroblasts   ATAC
WT_K1   WTFibroblasts   ATAC
WT_K2   WTFibroblasts   ATAC
WT_K3   WTFibroblasts   ATAC
```

```
#2023-June
#usage: Rscript sampletable_path Htseqcount_with_header_path
        ComparisonName NameOfTreatedSample
#remember to modify the ref level before running the script.

#silence the screen output when loading R packages, not supressing
        other output messages from other commands
suppressMessages(library(tidyverse))
suppressMessages(library(DESeq2))

#load packages
library(tidyverse)
library(DESeq2)
#read the input for R script
args <- commandArgs(trailingOnly = TRUE)
#sample info
print(args[1])
#count matrix
print(args[2])
#name for the plot
print(args[3])
#colnames for the DA peak list
print(args[4])

plotname <-args[3]
print(plotname)

#save the screen output into
sink(paste0(plotname,"DESeq2.R.out"))

sampletable <- read.table(args[1],sep="\t",header =TRUE)
sampletable$condition<-as.factor(sampletable$condition)

message("sampletable")
sampletable

message("genecount")
genecount <- read.table(args[2], header=TRUE,sep="\t",row.names=1)
head(genecount)
summary(genecount)

dds<-DESeqDataSetFromMatrix(countData=genecount, colData=sampletable,
        design = ~condition)

#define the reference level
dds$condition <- relevel(dds$condition, ref = "WTFibroblasts")
dds$condition

#testing different cutoffs to remove peaks woth low readcount from all
        samples
summary(rowSums(counts(dds)) >= 18)
summary(rowSums(counts(dds)) >= 30)
summary(rowSums(counts(dds)) >= 36)
summary(rowSums(counts(dds)) >= 40)

#filtering the peaks with low read count, total number of reads from
        all samples less than 40
keep <- rowSums(counts(dds)) >= 40
dds<-dds[keep,]
```

```r
#perform normalization (with the effective library size) and
        differnetial analysis
message("DA analysis")
dds<-DESeq(dds)

#resultsNames(dds)

head(results(dds, alpha=0.05))
summary(results(dds, alpha=0.05))
summary(results(dds, alpha=0.01))
summary(results(dds, alpha=0.001))

#transform the results
resLFC <- lfcShrink(dds, coef=2, type="apeglm")
resLFCOrdered <- resLFC[order(resLFC$pvalue),]
DEresult<-as.data.frame(resLFCOrdered)
DEresult$Peak<-row.names(DEresult); row.names(DEresult)<-NULL

#produce and save MA and volcano plots
#get the range of log2FC
myy<-max(round(abs(DEresult$log2FoldChange)))

#vocalno
p1<-ggplot(DEresult) +geom_point(aes(x=log2FoldChange,y=(-
        log10(padj)),color=(padj<0.05)),size=0.5) +
    xlim(-myy,myy) + theme_bw() +
    scale_color_manual(values=c("grey","blue")) +
    theme(axis.text=element_text(size=12,colour = "black"),
        axis.title=element_text(size=14)) +
    ggtitle(paste0(plotname,"ATACseq DESeq2\nVolcano plot of
        diffenrentialy gene expression")) +
    labs(x=paste0("log2FC",args[3]))

#MAplot
p2<-ggplot(DEresult) + geom_point(aes(x=baseMean, y=log2FoldChange,
        color=(padj<0.05)),size=0.5) +
    scale_x_log10() + ylim(-myy,myy) + theme_bw() +
    scale_color_manual(values=c("grey","blue")) +
    theme(axis.text=element_text(size=12,colour =
        "black"),axis.title=element_text(size=14)) +
    ggtitle(paste0(plotname," ATACseq DESeq2\n MA plot of DA peaks"))
        +
    labs(x="Normalized readcount")

pdf(paste0("ATAC.",plotname,"volcano_MAplot.pdf"),width=7,height=6)
p1
p2
dev.off()
sink()


#save the results
message("save the DA results")
myname<-args[4]
message(paste0("saving results for",myname))
colnames(DEresult)<-c(paste0(myname,"_baseMean"),
        paste0(myname,"_log2FC"), paste0(myname,"_lfcSE"),
        paste0(myname,"pval"), paste0(myname,"padj"),"Peak")
write.table(DEresult,file=paste0("DESeq2",plotname,".resLFCOrdered.table"),quote=F,col.names=T,row.na
message("save the R enviroment")
save.image(paste0("DESeq2",plotname,".Rdata"))
```

### Differential analysis of G4 ChIpseq data with/out CNV correction via DiffBind

#### pre-peak calling

read trimming, alignment and filtering are the same as decribed in ATACseq data processing

#### peak calling and IDR filtering

##### preparing the bamfiles

```
# for each biological replicate of a sample, there are 3 technical
        replicates for IP samples and one Input sample
WT_rep1.IP1.bam
WT_rep1.IP2.bam
WT_rep1.IP3.bam
WT_rep1.Input.bam

WT_rep2.IP1.bam
WT_rep2.IP2.bam
WT_rep2.IP3.bam
WT_rep2.Input.bam

BS_rep1.IP1.bam
BS_rep1.IP2.bam
BS_rep1.IP3.bam
BS_rep1.Input.bam

BS_rep2.IP1.bam
BS_rep2.IP2.bam
BS_rep2.IP3.bam
BS_rep2.Input.bam

# subsample 30Mil reads from each IP samples and merge them
WT_rep1.merge.bam
WT_rep2.merge.bam
BS_rep1.merge.bam
BS_rep2.merge.bam

# merge the input samples as well so that during peak calling, macs2
        will not downsample the IP samples as there are less reads in
        Input smaples
WT_input.bam
BS_input.bam
```

##### peak calling with macs2 for files merged from technical replicates

```
#!/bin/bash
#usage: scrit.sh IP_file Input_file
file=$1
fbname=$(basename $file .bam)
echo $fbname

#the control file for peak calling
file2=$2
echo $file2
macs2 callpeak --tempdir /tmp/dsu/ -g hs \
    -t $file -c $file2 \
    --outdir /tmp/dsu/ \
    -n "$fbname" --seed 11 --cutoff-analysis --keep-dup all \
    2> "$fbname".callpeak.log
```

##### IDR filtering on the peaks called from 2 biological replicates

```
#!/bin/bash
file1=$1;
echo $file1
fbname1=$(basename $file1 _peaks.narrowPeak )
echo $fbname1

file2=$2;
echo $file2
fbname2=$(basename $file2 _peaks.narrowPeak )
echo $fbname2
#sort the narrowPeak files by the Signal value, which is the default
        setting
    sort -k7,7nr $file1 >"$file1".Signal
    sort -k7,7nr $file2 >"$file2".Signal
    idr --samples "$file1".Signal "$file2".Signal --rank signal.value
        --input-file-type narrowPeak --log-output-file
        IDR_"$fbname1"_"$fbname2".Signal.idr_log  --plot --output-file
        IDR_"$fbname1"_"$fbname2".Signal.idr

#filter the peaks with the threshold of IDR<0.05
# remove peaks not on cannonical chromosomes
# remove peaks on sex chromosomes
cat IDR_WT.Signal.idr| awk '{if($5 >= 540) print $0}' |awk '$1!~/_/'
        |grep -v chrX|grep -v chrY |awk '{print
        $1"\t"$2"\t"$3}'>IDR_WT.Signal_IDR0.05.bed

cat IDR_BS.Signal.idr| awk '{if($5 >= 540) print $0}' |awk '$1!~/_/'
        |grep -v chrX|grep -v chrY |awk '{print
        $1"\t"$2"\t"$3}'>IDR_WT.Signal_IDR0.05.bed
```

#### making greylist for later differential analysis

##### calling peaks with input sample

```
#calling narrowPeaks with -f BAM
macs2 callpeak --tempdir /tmp/dsu/ -g hs \
    -t $Input_file \
    --outdir /tmp/dsu/ \
    -n "$fbname"_BAM \
    --seed 25 -B --SPMR --keep-dup all

macs2 callpeak --tempdir /tmp/dsu/ -g hs \
    -t $Input_file \
    --outdir /tmp/dsu/ \
    -n "$fbname"_BAM_broad \
    --seed 25 -B --SPMR --keep-dup all --broad

macs2 callpeak --tempdir /tmp/dsu/ -g hs \
    -t $Input_file \
    --outdir /tmp/dsu/ \
    -n "$fbname"_BAMPE \
    --seed 25 -B --SPMR --keep-dup all -f BAMPE

# for each input sample, merge the all the above peaks called from
        each command
cat WT_input_peaks.narrowPeak WT_input*_peaks.broadPeak |awk '$9>=3'|
        awk '{print $1"\t"$2"\t"$3"\t"$4}' |sort -k1,1 -k2,2n
        |bedtools merge -i stdin
        >WT_input.merge_broad_narrow_peaksq0.001.bed

cat BS_input_peaks.narrowPeak BS_input*_peaks.broadPeak |awk '$9>=3'|
        awk '{print $1"\t"$2"\t"$3"\t"$4}' |sort -k1,1 -k2,2n
        |bedtools merge -i stdin
        >BS_input.merge_broad_narrow_peaksq0.001.bed
```

###### generating greylist via DiffBind

###### here it is not the differential anlaysis. It is to generate a list of regions that have high coverage in the input samples

- folowing DiffBind vegnette, load data into DiffBind and proceed until the step of differntial anlaysis
- extract and save the greylist in bed format as `DiffBind_greylist.bed`

###### preparing the final greylist for Differential analysis

```
cat WT_input.merge_broad_narrow_peaksq0.001.bed
        BS_input.merge_broad_narrow_peaksq0.001.bed
        DiffBind_greylist.bed|sort -k1,1 -k2,2n |bedtools merge -i
        stdin |cut -f1-3 >Greylist.bed
```

#### Differential analysis in DiffBind

###### prepare metadata for samples in the differential analysis

note that here technical replicates of the same biological replicates are treated as individual samples instead of being collasped as one sample. Metadata is stored as `` ### Differential analysis in R with DiffBind

###### *read in the data and count reads in peaks*

```
#an example: Rscript ../DiffBind.R Sample.info BSvsWT Greylist.bed

#silence the screen output when loading R packages, not suppressing
        other output messages from other commands
#suppressMessages(library(DiffBind))
#load packages
library(parallel)
library(DiffBind)
library(BiocParallel)
register(MulticoreParam(workers = 6))

#read the input for R script
args <- commandArgs(trailingOnly = TRUE)

#sample metadata
message("sample sheet is", args[1])
sampleinfo<-args[1]

sample <-read.table(sampleinfo,sep="\t",header=T)

#name of plots
message("mysample name is ",args[2])
mysample<-args[2]

#greylist
#greylist is not essential. Greylist can be generated internally by
        DiffBind when gDNA data is included

mygreylist<-read.table(args[3],header=FALSE,sep="\t")
colnames(mygreylist) <- c("chr","start","end")
mygreylist<-GRanges(mygreylist)

sink(paste0(mysample,"Diffbind.R.out"))
#Read in the samples and their bamfiles
#strangely R encounter erros when I Run the script.
#when I enter R and read in the sampleinfo, things are fine
myG4 <- dba(sampleSheet=sample)
myG4

#remove regions in the blacklist,#remove regions in the greylist
```

```
myG4 <- dba.blacklist(myG4,blacklist=DBA_BLACKLIST_HG38,greylist=mygreylist)

#count the reads in peaks!
#crtitial to set Summits=FALSE!!!!
message("\ncount reads in peaks...")
myG4<-dba.count(myG4,summits=FALSE)
```

***optional copy number correction:***

- export the intervals of merged peaks in myG4 object
- assign the scaling factor to those intervals the same way as it in ATACseq data
- output: `scaling_factors.bed` with 4 columns, chr start end CNR

```
awk -v OFS='\t' '{print $1,$2,$3,$6}' scaling_factors.bed|awk -v
        OFS='\t' '{if ($4 >1) print $1,$2,$3,$4,1;else print
        $1,$2,$3,1,1/$4}'|sort -k1,1 -k2,2n > myscale

#read myscale into R
#the bed intervals are in the same order as the count matrix stored in
        myG4 object
#modify the readcount

#myG4$peaks[[1]] --> the first sample
#[,c(4,5,6,7)] --> 4,5,6,7 column for the first sample, which are the
        read count infomation in peaks

#in my case the 1-5 samples are BS and 6-11 samples are WT. Modify the
        count as the following
myG4$peaks[[1]][,c(4,5,6,7,8)]<-myG4$peaks[[1]]
        [,c(4,5,6,7,8)]/myscale[,4]
myG4$peaks[[2]][,c(4,5,6,7,8)]<-myG4$peaks[[2]]
        [,c(4,5,6,7,8)]/myscale[,4]
myG4$peaks[[3]][,c(4,5,6,7,8)]<-myG4$peaks[[3]]
        [,c(4,5,6,7,8)]/myscale[,4]
myG4$peaks[[4]][,c(4,5,6,7,8)]<-myG4$peaks[[4]]
        [,c(4,5,6,7,8)]/myscale[,4]
myG4$peaks[[5]][,c(4,5,6,7,8)]<-myG4$peaks[[5]]
        [,c(4,5,6,7,8)]/myscale[,4]

#Correcting the read count for WT
myG4$peaks[[6]][,c(4,5,6,7,8)]<-myG4$peaks[[6]]
        [,c(4,5,6,7,8)]/myscale[,5]
myG4$peaks[[7]][,c(4,5,6,7,8)]<-myG4$peaks[[7]]
        [,c(4,5,6,7,8)]/myscale[,5]
myG4$peaks[[8]][,c(4,5,6,7,8)]<-myG4$peaks[[8]]
        [,c(4,5,6,7,8)]/myscale[,5]
myG4$peaks[[9]][,c(4,5,6,7,8)]<-myG4$peaks[[9]]
        [,c(4,5,6,7,8)]/myscale[,5]
myG4$peaks[[10]][,c(4,5,6,7,8)]<-myG4$peaks[[10]]
        [,c(4,5,6,7,8)]/myscale[,5]
myG4$peaks[[11]][,c(4,5,6,7,8)]<-myG4$peaks[[11]]
        [,c(4,5,6,7,8)]/myscale[,5]

#Round the read count to integer numbers
myG4$peaks[[1]][,6]<-round(myG4$peaks[[1]][,6])
myG4$peaks[[2]][,6]<-round(myG4$peaks[[2]][,6])
myG4$peaks[[3]][,6]<-round(myG4$peaks[[3]][,6])
myG4$peaks[[4]][,6]<-round(myG4$peaks[[4]][,6])
myG4$peaks[[5]][,6]<-round(myG4$peaks[[5]][,6])
myG4$peaks[[6]][,6]<-round(myG4$peaks[[6]][,6])
myG4$peaks[[7]][,6]<-round(myG4$peaks[[7]][,6])
myG4$peaks[[8]][,6]<-round(myG4$peaks[[8]][,6])
myG4$peaks[[9]][,6]<-round(myG4$peaks[[9]][,6])
myG4$peaks[[10]][,6]<-round(myG4$peaks[[10]][,6])
myG4$peaks[[11]][,6]<-round(myG4$peaks[[11]][,6])
```

***Data normalization and differential analysis***

```
myG4<-dba.normalize(myG4,normalize=DBA_NORM_NATIVE,method=DBA_ALL_METHODS,background=F,l
        DBA_LIBSIZE_PEAKREADS)

myG4 <-dba.contrast(myG4,reorderMeta=list(Factor="WT"))
myG4<-dba.analyze(myG4,method=DBA_ALL_METHODS)
dba.show(myG4, bContrasts=TRUE)

#save the MA plot
pdf(paste0(mysample,"DiffBind_countreads_in_peak.MAplot.pdf"))
dba.plotMA(myG4, method=DBA_DESEQ2, sub=paste0(mysample," DESeq2,reads
        in peak"))
dba.plotMA(myG4, method=DBA_DESEQ2, sub=paste0(mysample," DESeq2,reads
        in peak"),yrange=c(-4,4))
dba.plotMA(myG4, method=DBA_EDGER, sub=paste0(mysample," edgeR,reads
        in peak"))
dba.plotMA(myG4, method=DBA_EDGER, sub=paste0(mysample," edgeR,reads
        in peak"),yrange=c(-4,4))
dev.off()

#write out the results
message("Save results with edgeR normalization ")
#extract the report
myG4_edgeR<-dba.report(myG4,th=1,method=DBA_EDGER)
mytable <-data.frame(
    seqnames = seqnames(myG4_edgeR),
    starts = start(myG4_edgeR) - 1,
    ends = end(myG4_edgeR),
    Conc = elementMetadata(myG4_edgeR)$Conc,
    BSConc = elementMetadata(myG4_edgeR)$Conc_BS,
    WTConc = elementMetadata(myG4_edgeR)$Conc_WT,
    log2FC = elementMetadata(myG4_edgeR)$Fold,
    pvalue = elementMetadata(myG4_edgeR)$"p-value",
    FDR = elementMetadata(myG4_edgeR)$FDR
  )

write.table(mytable,
  paste0(mysample,"_Diffbind.RiP_edgeRnorm.DAresults.bed"),
  col.names = F,
  quote = F,
  row.names = F,
  sep = "\t"
)
```
